# Single-molecule detection and super-resolution imaging with a portable and adaptable 3D-printed microscopy platform (Brick-MIC)

**DOI:** 10.1101/2023.12.29.573596

**Authors:** Gabriel G. Moya Muñoz, Oliver Brix, Philipp Klocke, Paul D. Harris, Jorge R. Luna Piedra, Nicolas D. Wendler, Eitan Lerner, Niels Zijlstra, Thorben Cordes

## Abstract

Over the past decades, single-molecule and super-resolution microscopy have advanced and represent essential tools for life science research. There is,however, a growing gap between the state-of-the-art and what is accessible to biologists, biochemists, medical researchers or labs with financial constraints. To bridge this gap, we introduce Brick-MIC, a versatile and affordable open-source 3D-printed micro-spectroscopy and imaging platform. Brick-MIC enables the integration of various fluorescence imaging techniques with single-molecule resolution within a single platform and exchange between different modalities within minutes. We here present variants of Brick-MIC that facilitate single-molecule fluorescence detection, fluorescence correlation spectroscopy and super-resolution imaging (STORM and PAINT). Detailed descriptions of the hardware and software components, as well as data analysis routines are provided, to allow non-optics specialist to operate their own Brick-MIC with minimal effort and investments. We foresee that our affordable, flexible, and open-source Brick-MIC platform will be a valuable tool for many laboratories worldwide.

## Introduction

Research in the molecular life sciences, biomedicine and under clinical settings heavily relies on the use of light microscopy(*1, 2*), biophysical techniques(*3, 4*), spectroscopic assays, e.g., PCR(*5*), ELISA(*6*), DNA sequencing(*7–9*) and many others(*10–12*). There is rapid advancement of both the instrumentation and the assays to achieve better spatial- and temporal-resolution or higher sensitivity for various applications including virus detection(*13– 17*). This progress, however, typically takes place in engineering and (bio)physics labs and it is difficult to benefit from it in applied- or industry research and under clinical settings(*1, 18*). Reasons for this can be the mere dimensions of a setup, or the inability to operate it outside of controlled lab conditions, e.g., due to missing temperature control or lack of mechanical stability. Thus, many advanced techniques cannot be used in high biosafety labs, on field trips, research ships, in hospitals, doctor’s practices, and other locations outside the lab. Consequently, there is a growing gap between the possibilities of the state-of-the-art in microscopy and spectroscopy and what is accessible to all interested users(*18, 19*). While there are core facilities for imaging and biophysical techniques, these remain too few, might only provide limited infrequent access or come with the requirement for travel. All this poses fundamental limitations since many biological and medicinal studies require long iterative refinement, samples may have to be studied locally and point-of-care applications using advanced techniques are simply not feasible(*18, 19*).

In recent years, different research groups and companies have started to bridge this gap by miniaturizing microscopy research platforms and reducing costs of commercial systems. Currently, the available compact microscopy setups offer high spatial- and temporal-resolution or high sensitivity. The setups often utilize commercially available optomechanical components, as has been implemented in the smfBOX(*20*) and miCUBE(*21*). On one hand, despite their performance, these setups cannot be used easily outside of optical laboratories and require substantial expertise for setting them up, maintaining them, and operating them – making them less suitable for application-oriented users. On the other hand, Oxford Nanoimager’s video-based device is small, powerful, and user-friendly, but is inflexible in terms of microscope modalities(*22–24*). Another drawback of all the aforementioned microscopes are high costs, which are well over 100,000 €. On the contrary, 3D-printing with plastic materials has gained popularity to replace expensive optomechanics and parts of the microscope frame. AttoBright, a user-friendly, minimalist, confocal microscope, showed that a 3D-printed setup can facilitate single-molecule detection(*25*). It comes, however, with substantial limitations in terms of data quality and general adaptability compared to the smfBOX or miCUBE. Another option, the “UC2”, a camera-based microscope, represents an adaptable platform designed for educational purposes, which allows the realization of various imaging modalities, including bright-field, dark-field, fluorescence microscopy(*26*) and most recently single-molecule localization microscopy(*27*).

To overcome these limitations, we here introduce an open-source microscopy platform called Brick-MIC, which uses a combination a modular 3D-printed scaffold with a minimal number of optical components. It offers the possibility to realize a variety of (fluorescence) microscopy modalities, e.g., confocal and video detection, fluorescence spectroscopy assays, state-of-the-art single-molecule detection and super-resolution optical imaging using the same platform. The scaffold of the microscope consists of four layers made from 3D-printed plastic material (Figure 1): a sample holder, an excitation layer, a detection layer, and a base plate for the interchangeable excitation and detection layers. All technical drawings and detailed descriptions on how to build and assemble the different Brick-MIC modalities are available with this manuscript. We further provide (compiled) python-based data acquisition and analysis software to perform experiments with different confocal modalities, making the Brick-MIC a true open-source microscopy platform. The only requirement to start your own Brick-MIC is access to a standard, low-cost 3D printer and to purchase of a minimal list of opto-mechanical and optical components.

**Figure 1.**
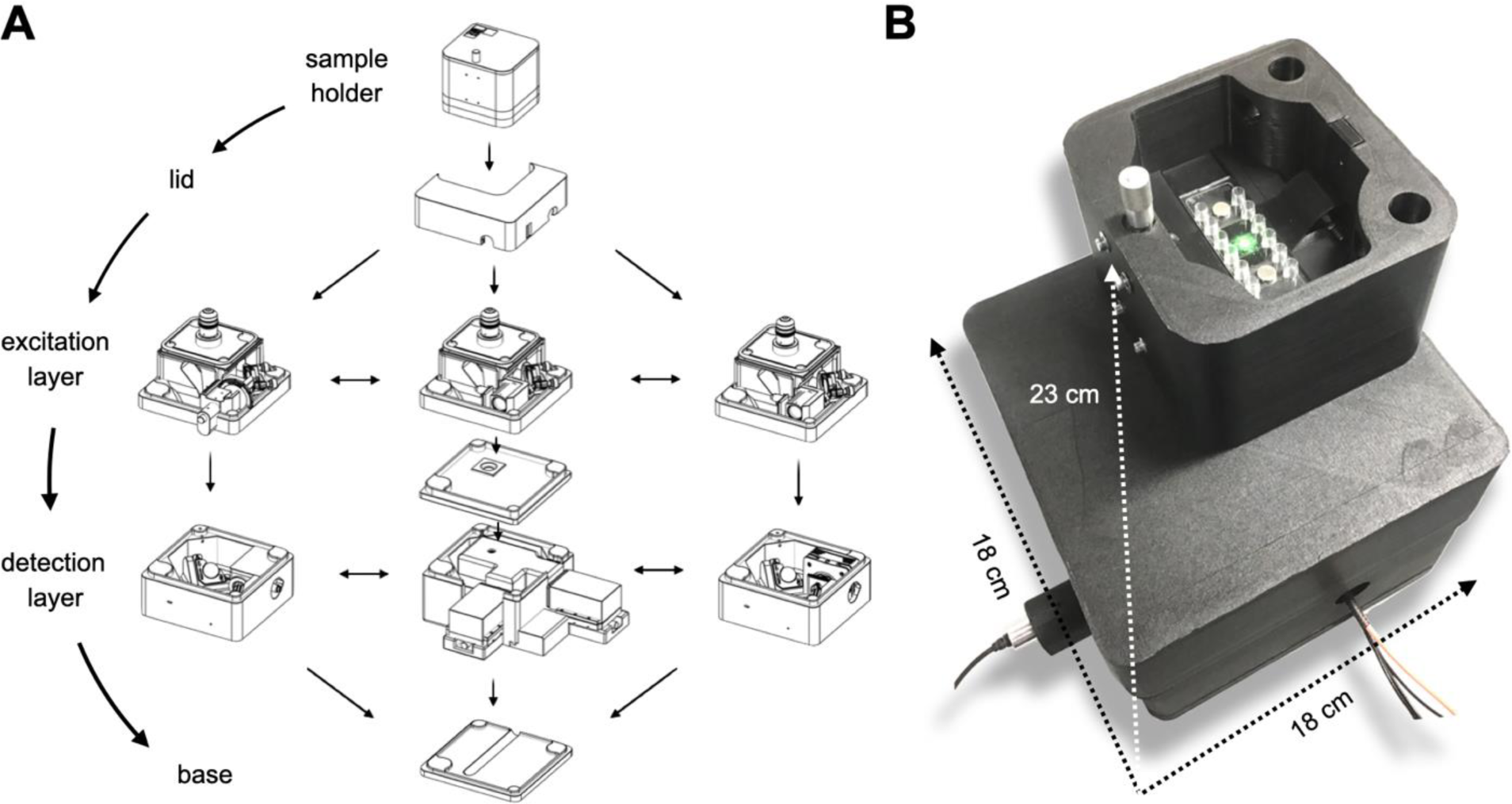
Overview of the adaptable Brick-MIC design. A) The platform uses shape-complementary parts that can be stacked (Lego-like or Japanese “poka yoke”), consisting of different layers: the sample holder, lid, excitation, detection and base layers. The excitation and detection layers are interchangeable, allowing to establish different imaging methods with the same platform and easy exchange of components. B) Photograph of a confocal Brick-MIC modality as used below for analysis of labelled biomolecules in free diffusion (section “Single particle fluorescence detection and fluorescence correlation spectroscopy”).

To date, we have established various distinct modalities of the platform, from which we use three as representative examples in this paper. The first is a confocal microscope for single particle detection, fluorescence correlation spectroscopy (FCS(*28*)) and time-correlated single-photon counting (TCPSC(*29, 30*)), with which we were able to detect individual fluorescent particles, such as freely-diffusing nano-sized fluorescent beads, dye molecules, and labelled bio-molecules of varying sizes. Secondly, we established a two-color confocal microscope that we used for single-molecule Förster resonance energy transfer (smFRET(*31, 32*)) experiments with microsecond alternating-laser excitation (µsALEX(*33, 34*)). Finally, we realized a camera-based microscope that allows standard widefield or darkfield imaging and can be upgraded to an epi-fluorescence microscope for fluorescence and super-resolution imaging via STORM(*35, 36*) and DNA-PAINT(*37, 38*). We consider our approach a “Swiss-knife” microscope with full flexibility and hope that Brick-MIC will be used inside and outside of research laboratories in the future due to its small size and portability, high stability and state-of-the-art performance.

## Results

### General considerations for the design of the Brick-MIC platform

Brick-MIC was designed with the philosophy to create a user-friendly, portable and stable, cost-effective and adaptable platform that can be used outside of optical labs under ambient light. To reduce costs and allow robust operation, we established an adaptable microscope body which is fully 3D-printed. This platform is combined with optical components such as mirrors, filters, etc. from commercial suppliers to establish one microscope modality. The printing templates (https://zenodo.org/records/10441063) and a list of optical and optomechanical components are provided as Supplementary Information (Brick-MIC component list). Once the microscope frames for all modalities are printed, the Brick-MIC platform enables rapid exchange between distinct microscopy modalities within minutes. We demonstrate both the straightforward assembly and the rapid exchange of modalities in Supplementary Videos 1 ,2, and 3, respectively. The enclosed sample holder is suitable for the incorporation of microfluidics components, such as Ibidi microfluidics slides and tubing(*39*), and allows to fix microscope slides with magnets (Supplementary Video 1, 2). A single-axis translation stage, which is integrated into a rack and pinion system, controls the Z-axis position of the coverslip and sample, and ensures high positional stability (Figure 1). The design was tested to generally ensure mechanical stability via an accurate fit of the different layers and vibration dampening by use of thermoplastic Polyurethane (TPU), a resilient and rubber-like material, as the base layer of the microscope (Figure 1 and Supplementary Video 1). The detection layers of Brick-MIC are all equipped with essential optical components mounted onto piezo motors for convenient alignment of the setup and use of auto-calibration software (see below). Importantly, all modalities of Brick-MIC, presented in this paper require low investment costs between 10.000 and 30.000 €, can be operated with software acquisition packages provided by the respective detector suppliers or with publicly available software (see below), which do not require expensive software licenses, in line with open science practices(*19*).

### Single particle fluorescence detection and fluorescence correlation spectroscopy (FCS)

Confocal microscopy can be used for sensitive detection of (individual) fluorescently-labeled particles and molecules in free diffusion or flow, either via burst detection(*40*) or FCS(*28*). Based on the idea to use Brick-MIC outside of optical labs in biomedical research, clinical diagnostics, or environmental monitoring we established a basic modality with a single cw-laser excitation source and dual-channel detection using photomultipliers, PMTs in single-photon counting mode (Figure 2A/B). The confocal geometry of the setup is achieved using an inversely mounted parabolic collimator that couples the emitted light into an optical fiber (OF). The OF serves as a pinhole (PH) with a diameter related to its core size. The emitted light is captured by a high NA water-immersion microscope objective (60X, NA = 1.2, Olympus – UPLSAPO60XW) and directed by the OF into an external detection box. Here, the emission is spectrally separated via a dichroic mirror (DM) into short (green) and long wavelength emission (red) before reaching two photomultiplier tubes (PMTs; Figure 2B).

**Figure 2.**
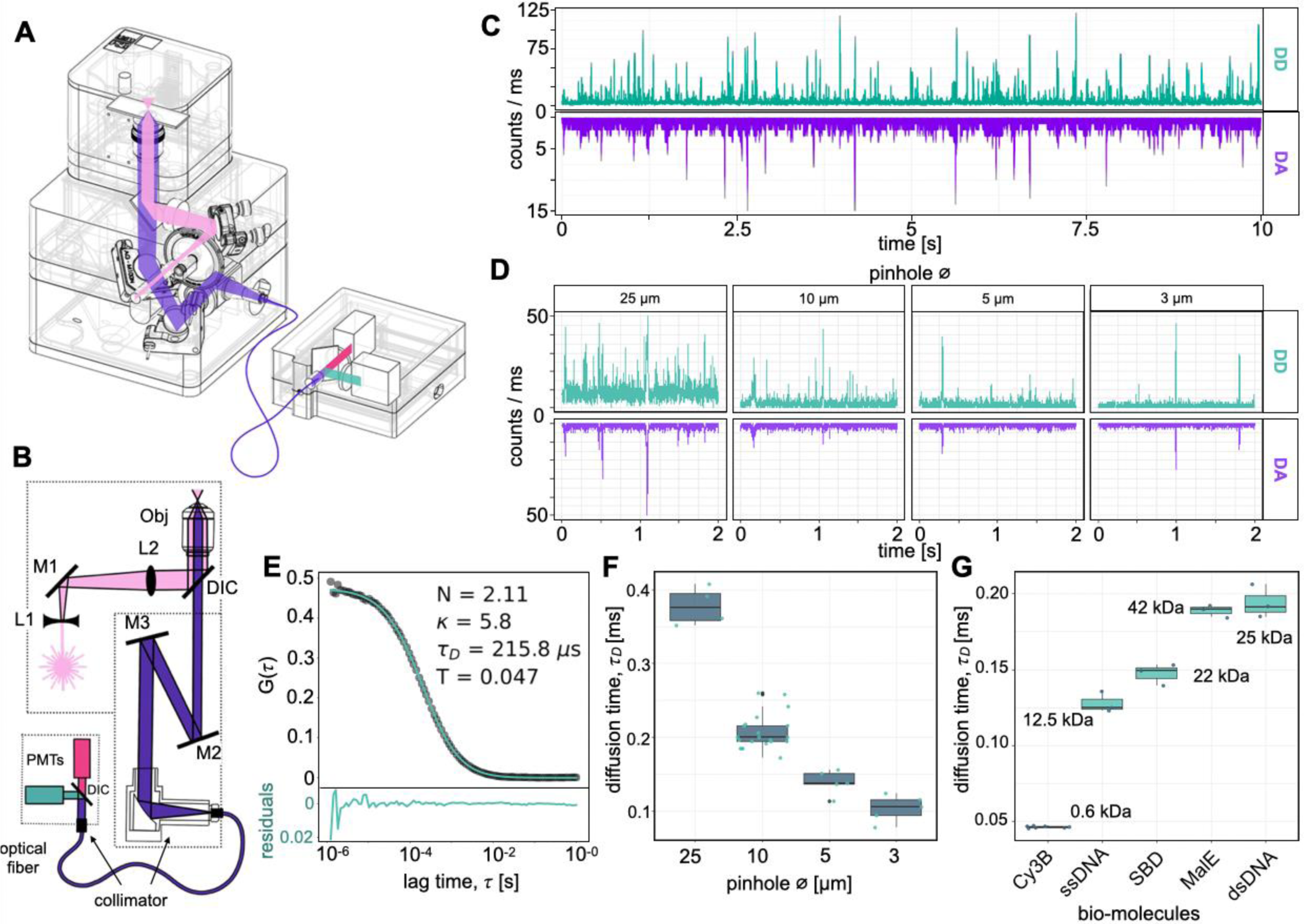
Single-molecule detection and fluorescence correlation spectroscopy (FCS) with Brick-MIC. **A)** CAD model showing Brick-MIC with optical components and light paths (pink: excitation; purple: emission; green and red: spectral split emission after dichroic mirror (DIC)). **B)** Optical layout: a single laser diode (pink) expanded by a telescope (bi-concave lens L1, plano-convex lens L2) focuses into the sample through an objective lens (Obj). Emission is collected and directed by mirrors on piezo-directed optical mounts (M2 and M3) through an inversely mounted reflective collimator, coupling the emission into an optical fiber (OF). The fiber directs light to an external detection box, spectrally separated by a dichroic mirror, detected by PMTs. **C)** Single molecule time trace of 100 pM dsDNA labeled with Cy3B (D) and ATTO647N (A) at a 13 bp inter-dye distance. Bursts recorded in donor (DD) and acceptor emission (DA) channels under continuous green excitation. **D)** Zoom-in on time traces (max counts: 50 kHz) of 500 nm Tetraspeck-beads in laminar flow (2.5 µl/s) with different optical fiber (OF) core diameters acting as a pinhole (PH). **E)** Representative FCS curve of a 5 nM 40-mer dsDNA sample labeled with Cy3B, using a 10 µm pinhole core diameter; fit parameters: number of molecules N, geometry parameter κ, diffusion time τD, and triplet-fraction T. **F)** Boxplot of FCS-based average diffusion times of a 5 nM 40-mer dsDNA sample labeled with Cy3B measured with different pinhole (PH) sizes. **G)** Boxplot of FCS-based average diffusion times of various biomacromolecules labeled with Cy3B with different masses and hydrodynamic radii, recorded using a 10 µm PH. Data acquired via NI-Card. Error bars: standard deviations from n > 3 repeats.

Signals of both PMTs are read out via an affordable single-photon counter (MCC-DAQ-USB-counter)(*25*) in combination with a python software provided with this manuscript. The data can be used for online inspection of fluorescence time traces in binned format, e.g. for calibration purposes, or can be exported to hdf5 single-photon data (https://github.com/harripd/mcc-daq-acquisition) for further processing. Importantly, alignment of the setup can be achieved with an automated self-alignment procedure via two piezo mirrors using concentrated solutions of e.g., 100 nM Cy3B (see Online Methods and Supplementary Video 4). The self-alignment procedure was directly implemented into the aforementioned Python code and allows exchange of modalities within minutes (see Supplementary Video 3 and 4). A notable benefit of this design is the convenient accessibility and interchangeability of the OF, serving as a PH with tunable diameter. This allows for easy modifications to the detection volume of the microscope by selecting optical fibers with varying core diameters, thereby offering enhanced flexibility in tailoring the parameters to specific experimental requirements.

Using this setup, we observed single fluorescently-labeled nanoparticles of varying diameters ≥100 nm (Supplementary Video 2) and individual fluorophore-labeled double-stranded DNA (dsDNA) molecules. Diffusional transits of donor-acceptor labeled dsDNA molecules (donor Cy3B, acceptor ATTO647N in 13 bp distance) were clearly visible as coinciding bursts in both detection channels (Figure 2C). Variation of the OF core size reveals the effect of shrinking excitation volume in fluorescence time traces (Figure 2D). Larger PH sizes, i.e., large detection volumes, show higher and longer signal periods, higher burst detection frequency as well as increased background compared to smaller PH diameters (Figure 2D and Supplementary Figure 2).

The high sensitivity of the setup suggests that it can readily be used for FCS measurements. We first tested and quantified parameters of a 10 nM solution of 40mer-dsDNA labeled with Cy3B (Figure 2E). For data acquisition we used either the described MCC-DAQ-USB-counter(*25*) or an NI-Card as described previously by Gebhard et al.(*41*) For FCS data analyses, we provide a jupyter notebook (https://github.com/PSBlmu/FCS---analysis), which uses functions of the publicly-available FRET-bursts script from the Weiss lab(*41, 42*), for analysis of the obtained single-photon data in the hdf5-(MCC-DAQ-USB-counter) or binary-format (NI-Card). With this procedure, we could extract molecular brightness B, diffusion time, triplet lifetime and their associated amplitudes from a standard two-component fit with diffusion and triplet (Figure 2D). The average diffusion times of dsDNA were ∼100, ∼150, and ∼200 µs for increasing PH diameters. The use of an OF with a core size diameter of 10 µm resulted in comparable results to those obtained from a custom-built confocal microscope(*41, 42*) which employs a 50 µm PH (Figure 2F and Supplementary Figure 3). The brightness of Cy3B was lower in the Brick-MIC microscope (10 kHz vs. 80 kHz per molecule, Supplementary Figure 3 and 4) either due to a reduced PH diameter or lower quantum efficiency of the PMTs. The analysis of diffusion times of biomolecules with varying masses and hydrodynamic radii revealed that both setups correctly assess the expected trends related to molecular mass differences (except for the non-spherical dsDNA sample; Figure 2G).

To demonstrate the flexibility of Brick-MIC, e.g., to use pulsed instead of continuous laser excitation, we exchanged the excitation layer to contain a reflective collimator as output of a fiber-coupled laser (Figure 3A/B and Supplementary video 3). As a proof-of-concept for TCSPC, the fluorescence lifetimes of Cy3B, Alexa546 and Atto550 fluorophore-labeled dsDNA samples were determined using a modified excitation layer on top of the emission layer of the FCS modality (Figure 2). We found lifetimes for Cy3B, Alexa546 and Atto550 on dsDNA of 2.82±0.01 ns, 3.91±0.01 ns and 4.15±0.04 ns (Supplementary Figure 5,6 and 7), respectively, matching values from a home-built cuvette-based setup with one APD (Supplementary 5,6 and 7), which was described before(*41*). Each of the triplicate measurements were done via pulsed laser excitation at 532 nm wavelength at a rate of 20 MHz and power of 55 µW (LDH-P-FA-530B with PDL 828 “Sepia II” controller, Picoquant) detected by a Multiharp 150 8N (Picoquant) for the PMT variant. For the APD variant we used a supercontinuum white light laser source (NKT SuperK Extreme EXW-12, NKT Photonics) with 80 MHz and power of 55 µW for excitation and detection by a Hydraharp 400 (Picoquant) as described before(*41*).

**Figure 3.**
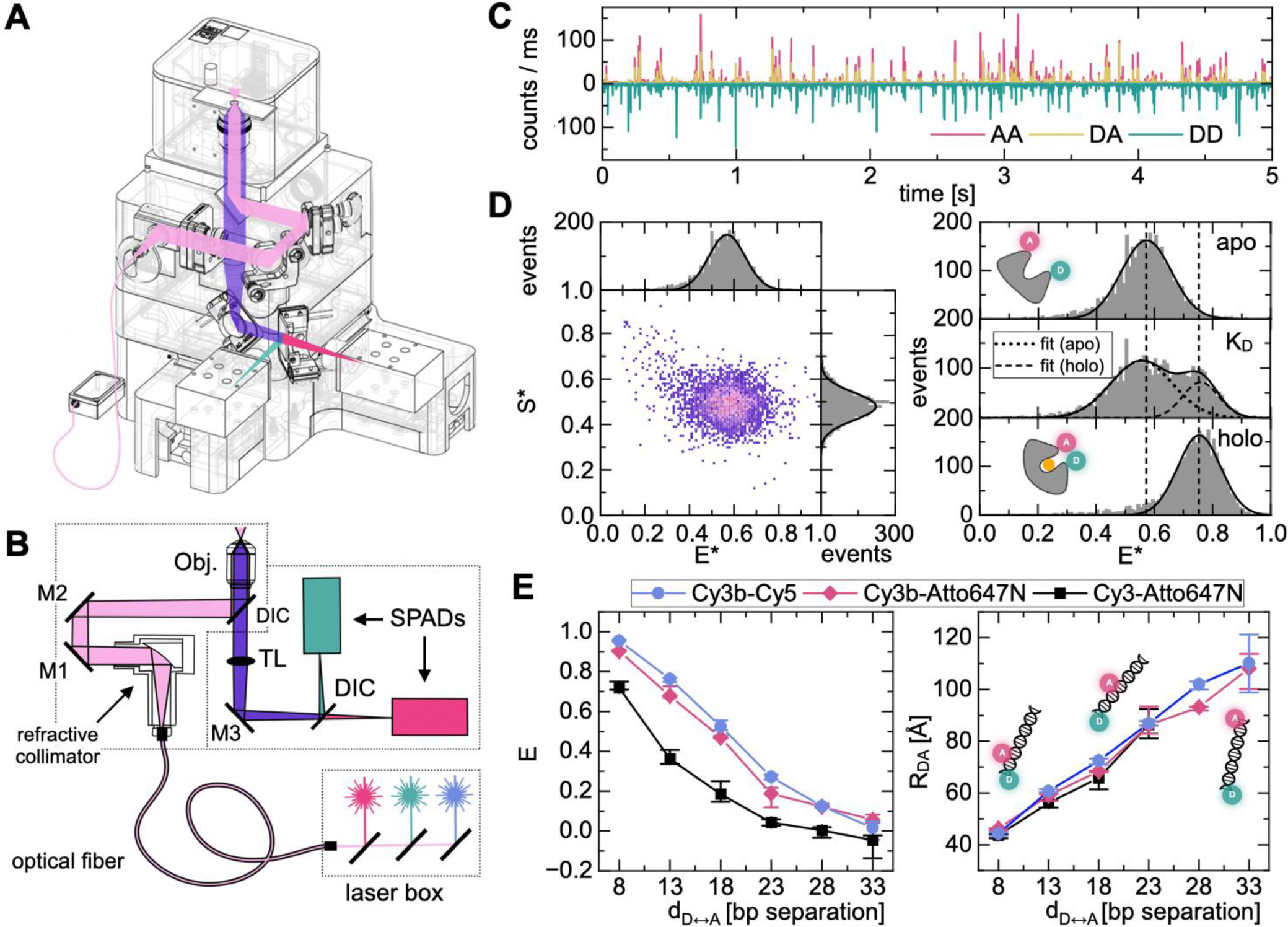
Single-molecule FRET and µsALEX modality. A) CAD model showing all optical components including the light path (pink: excitation; purple: emission; green and red: spectral split emission after the DIC). B) Overview of the optical layout: The modality uses an external laser coupled to the microscope through an OF. In this arrangement, the excitation beam is collimated with a reflective collimator and then focused into the sample through an objective lens. The emission is collected by the same objective lens and is further focused via an achromatic tube lens (TL) directly onto two different single-photon avalanche diodes (SPADs). Spectral separation is achieved by a DM and appropriate bandpass filters for each detector. The M3 Mirror and the DIC in the emission layer are mounted onto piezo-directed mirror holders, which are used to direct the photon stream into each detector channel. C) Time trace of a 100 pM dsDNA sample labeled with Cy3B donor (D) and ATTO647N acceptor (A) dye at a 13 bp inter-dye distance. Individual bursts are recorded in three different acquisition channels: DD (green) for donor excitation and donor emission; AA (red) for acceptor excitation and acceptor emission; and DA (yellow) for donor excitation and acceptor emission. D) Representative ES-histogram of the open conformation of the substrate binding protein MalE (left side). (Right side) Observation of different conformational states of MalE (open, closed, and KD conditions) for apo (no substrate), holo (100 µM maltose), and KD (1µM maltose). E) Determination of accurate FRET efficiency values and corresponding distances (RDA) using a DNA ladder for different dye combinations. Data obtained using acquisition via NI-Card (details see methods).

### Single-molecule FRET and ALEX

FRET and its single-pair equivalent spFRET (or smFRET) have become an established method in the prospering toolkit of integrative structural biology(*42–52*). With the smFRET method, it is possible to study biomacromolecules in aqueous solution at ambient temperature, and to identify conformational heterogeneity and sub-populations, measure accurate distances and to characterize conformational changes (kinetic exchange rates)(*32, 34, 41–47, 53–59*).

As expected from the sensitivity of the setup, it was possible to use the PMT-version of our microscope (shown in Figure 2) for ratiometric determination of apparent FRET efficiency E* of single donor-acceptor pairs under continuous green excitation with PMTs (Figure 2C). We note, however, that the sensitivity of the red detection channel was suboptimal and the available dynamic range of E* would be rather limited in comparison to our home-built setups. We thus further optimized the setup for this specific application. We thus improved the detection efficiency of the setup by use of two single-photon avalanche diodes (SPAD, model PDM 50-Micron, MPD) in the detection layer. These serve as single-photon counters and PHs simultaneously due to their small active detection area of 50 µm diameter^23^. As can be seen from the scheme of the optical setup (Figure 3A/B), the design greatly reduces the number of required optical and opto-mechanical components in comparison to standard home-built setups.

Employing this modality in combination with microsecond alternating laser excitation (µsALEX(*34*)) of fiber-coupled green and red lasers at an alternation rate of 20 kHz (50 µs excitation periods), we observed bursts from individual freely-diffusing FRET-labeled biomolecules (Figure 3C). The molecular brightness of Cy3B in this setup was around 60 kHz and thus far superior compared to the PMT variant (10 kHz) and only slightly lower than that of a home-built setup (80 kHz, Supplementary Figure 4). From these setups we extracted photon-streams relevant for ratiometric FRET determination (Figure 3D/E, apparent FRET efficiency E*) and apparent fraction of donor brightness (S*) for each single molecule transit through the confocal excitation volume. As shown for MalE, the periplasmic subunit of the maltose permease(*60*), both conformational states (apo: ligand-free open and holo: ligand-bound closed) and the ligand affinity can be characterized with single protein resolution(*55*) (Figure 3D and Supplementary Figure 16). These experiments utilize FRET as a qualitative indicator for conformational changes via E*, e.g., from low to high E* values, i.e., from long to short inter-dye distance, respectively.

We also assessed the ability of the setup to determine accurate FRET efficiencies, E, via correction of E* for all setup-dependent parameters, i.e., spectral crosstalk for donor leakage α, acceptor direct excitation δ, normalization of detection and quantum yield differences of acceptor and donor γ and normalization of excitation intensities and absorption cross sections of acceptor and donor dye ß, to obtain inter-dye distances, R_DA_, using established procedures(*42, 43, 47*). We compared the E values for an inter-dye base-pair (bp) separation of DNA ladder samples with different distances (in five bp steps) between the donor and acceptor dyes, ranging from 8 to 33 bp. The dependency of E as a function of bp separation differs for the three dye-pairs studied due to differing Förster distances (Figure 3E). We used values of R_0_ known from the literature: Cy3-ATTO647N: R_0_ = 5.1 nm(*56, 57*); Cy3B-ATTO647N: R_0_ = 6.7 nm(*57–59*); Cy3B-Cy5: R_0_ = 7.4 nm (for the latter see Online Methods). Importantly, all E values, which are independent of R_0_, are consistent between Brick-MIC and the corresponding experiments performed on a home-built confocal setup(*41, 42*) (Supplementary Figure 8-15). Furthermore, the derived R_DA_ values of all three dye-combinations are internally consistent and the inter-dye distances derived from the different dye pairs cannot be distinguished within the error margins, except for large distances (> 8 nm) that are outside of the sensitive dynamic range of the FRET approach (Supplementary Figure 8 and 9).

### Camera-based light microscopy, single-molecule fluorescence detection and super-resolution imaging

All our investigations show that confocal-based single-molecule detection is possible with a minimal and cost-effective 3D-printed microscope system (Figure 2/3). As a final step, we explored the potential of Brick-MIC for camera-based imaging. Figure 4 shows the developed wide-field epi-fluorescence modality. In this configuration, the excitation layer is linked to the light source via an OF, similar to the aforementioned µsALEX modality. We have, however, also tested the modality with a single laser pointer as was used for the confocal-based setup described in Figure 2.

**Figure 4.**
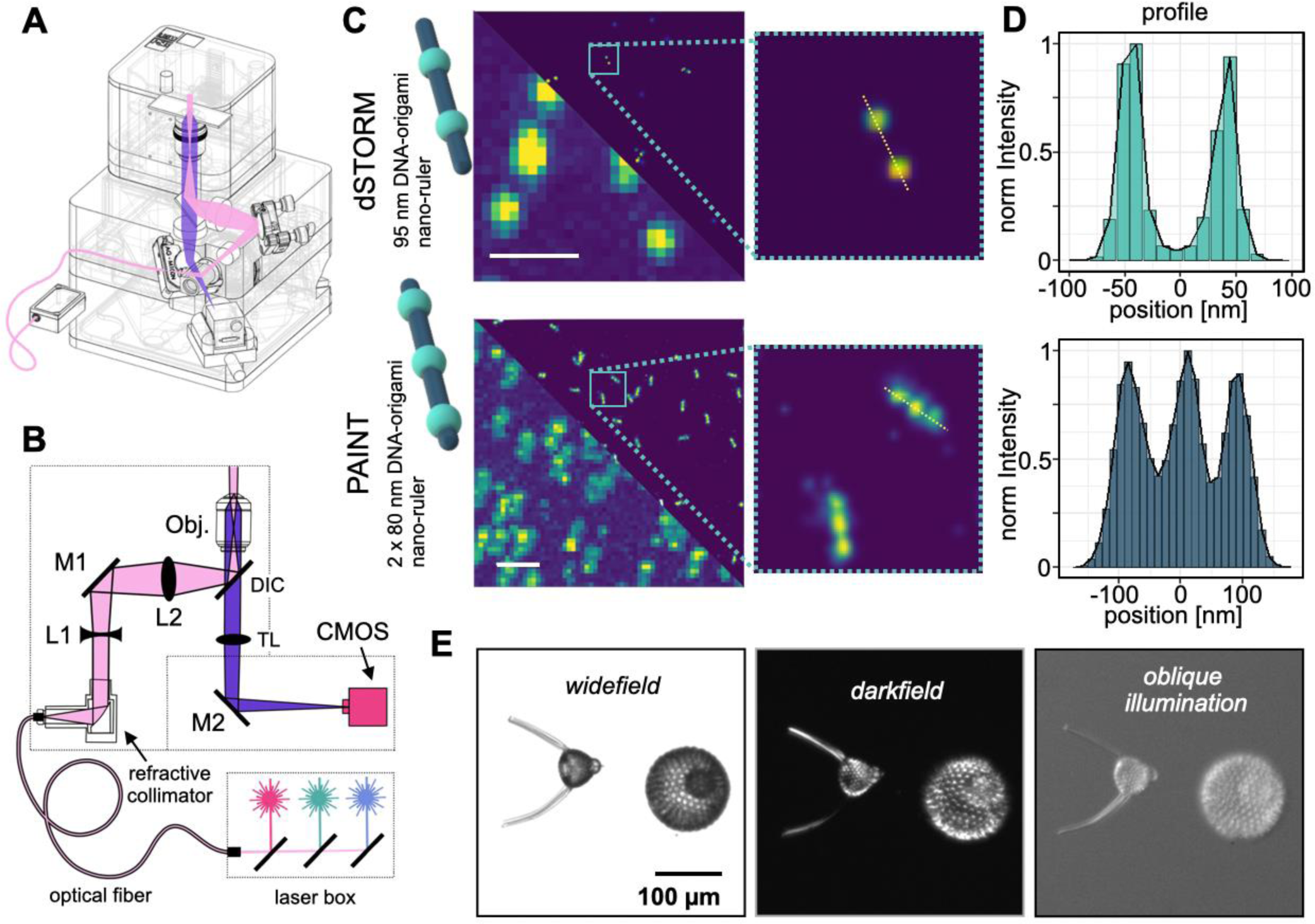
Camera-based light microscopy, single-molecule fluorescence detection and single molecule localization microscopy super-resolution imaging. A) CAD model showing all optical components, as well as the light path (pink: excitation; purple: emission). B) Overview of the optical layout: The modality uses an external laser box coupled to the microscope through an OF. In this arrangement, the excitation beam is collimated with a reflective collimator, which is expanded using a plano-concave lens (L1) and focused with a plano-convex lens (L2) into the back focal plane of the objective lens, resulting in an even illumination of the sample. The emission is collected by the same objective lens and focused via an achromatic tube lens (TL) onto a CMOS camera creating a real image of the sample. C) dSTORM and DNA-PAINT imaging (up and down, respectively) using 2 x 95 nm and 3 x 80 nm DNA-origami nano-ruler structures, respectively. Representative epi-fluorescence image and reconstructed super-resolution single-molecule localization image (left, white line represents 1 µm), with a zoom in (right). D) Representative cross-section profile of a single DNA-origami showing the zoom (yellow dotted line). E) Imaging of Radiolarians using classical contrast methods: transmission light microscopy (left), darkfield illumination (middle), and oblique illumination (right).

We then tested the microscope for localization-based super-resolution imaging using STORM(*35, 36*) and DNA-PAINT(*37, 38*) For this we obtained DNA-origami nano-rulers from GATTAquant with a single 95 nm distance (STORM) and two 80 nm distances (DNA-PAINT) between dye attachment positions on the respective origami structure(*54*). These fluorescent structures were sparsely immobilized on BSA/BSA-biotin coated surfaces and provided images as shown in the left panels of Figure 4C (epi-fluorescence, Supplementary Video 5/6). Applying thiol-containing photoswitching buffer(*35, 36*) or DNA-imager strands to the solution^33^ allowed using the blinking emission of individual labels to construct super-resolved images with the ImageJ plugin Thunderstorm(*61*). We found that beads were helpful, but not strictly necessary as fiducial markers to compensate for lateral drift, since lateral drift was < 250 nm over timespans of 30 mins (Supplementary Figure 18). With the setup both structures were resolvable, and we obtained a localization precision from isolated dyes or binding sites of ∼30 nm FWHM and 65 nm FWHM for STORM and PAINT, respectively (Figure 4D). Importantly, this localization precision was achieved from analyzing 2500 consecutive frames.

While the device was specifically tailored for fluorescence imaging, we also show its use in standard transmission light microscopy or contrast-enhancing techniques such as dark-field and oblique illumination, exemplified by images of Radiolarians recorded with the setup. To obtain these images, minimal modifications of the optical components were required. A LED desktop lamp was used as transmission light, positioned right on top of the sample holder. Additionally, the objective was interchanged for a 20x air objective with a low numerical aperture (NA=0.4, 1-U2B225, Olympus). Darkfield imaging, as well as oblique illumination, was achieved by covering the sample holder with aluminum foil and punching small holes in it with different patterns. For darkfield, a round pattern approximately 5 cm in diameter was used. For oblique illumination, a crescent moon shape was punctured in the foil approximately 3 cm away from the objective’s axis (Supplementary Figure 19).

### Discussion and Conclusion

As shown in this manuscript, Brick-MIC is a high-performance and cost-effective multi-functional microscopy platform that facilitates various state-of-the-art microscopy applications including single-molecule detection and super-resolution microscopy. The modular philosophy of the platform allows for development of additional new modalities not shown here, e.g., TIRF, FLIM or light-sheet microscopy. This versatility might make it a valuable technology platform not only for imaging, but also for a wide range of other scientific directions, e.g., flow cytometry or spectroscopy. Notably, the compact size of the platform offers advantages beyond the confines of the laboratory and that of previous works(*19–22, 24–26, 62*) in terms of performance and portability, as it enables fieldwork research, such as on-site water quality assays, or being deployed in restricted locations such as in high biosafety labs. The portability and small footprint of Brick-MIC make it ideal for conducting research in challenging conditions or real-world scenarios, opening new possibilities for scientific exploration.

The philosophy behind Brick-MIC was to optimize one technical modality (as shown above) while allowing flexibility to switch between different modalities through interchangeable modules. This principle allows to largely reduce the required components to build different microscopes. Particularly for the shown confocal modalities, the Brick-MIC design eliminated many components and allowed a simple construction for use of pinhole. In a conventional confocal setup the latter often involves a tube lens to focus the emission beam through the pinhole, followed by a second lens for focusing onto the detector. This part is also delicate in terms of alignment precision making it a weak point in terms of stability. In the Brick-MIC PMT variant, the entire pinhole assembly was streamlined into a single continuous unit. This unit comprises two collimators interconnected by an optical fiber, eliminating the need for alignment procedures between the components. For the APD detection variant, the entire assembly was simplified to a single lens by using the small aperture of the detectors as the pinhole (as also shown previously by Gambin and co-workers(*25*)). Consequently, the Brick-MIC alignment procedure could be simplified to a single variable, i.e., optimization of the first element before the pinhole allowing automated alignment via piezo-mirrors. Furthermore, the reduction in size significantly shortens the light path to approximately 30 cm, thereby enhancing the strength and stability of the Brick-MIC platform. As a result, despite the lower material stiffness of the printing filament PLA compared to steel (with a modulus of elasticity of around 3.5 GPa versus 200 GPa respectively), it proves suitable for state-of-the-art applications.We hope to have demonstrated here that the modularity of Brick-MIC allows for rapid prototyping, e.g., to ‘just quickly test of new configuration’, introduction of new parts (e.g., different sample holders), or to realize completely new ideas by creating modules within the platform without re-designing the setup completely. In that respect, we are currently testing our platform for applications in iSCAT microscopy (Brix, Moya, Cordes, Hartschuh, in preparation), but also consider replacing the PMT detection box simply by a fiber-coupled optical fluorescence spectrometer (Moya, Luna, Cordes, in preparation).

We believe that the scientific community and applied users in industry or biomedicine will also help us to further improve the technology by testing different optical components of distinct quality, e.g., low vs. high NA objectives. This could further help to balance the requirements for an affordable setup vs. performance. Additionally, extensions of the platform related to temperature control, an autofocus systems for video-microscopy and improved mechanical stability (by use of different 3D-printing materials) could be a topic of future research and engineering of the platform. Other directions would concern the establishment of detector modules where data recording and analysis are more user-friendly, e.g., by use of an action camera or a mobile phone, where data recording and analysis are even easier than demonstrated here.

## Online Methods

### Sample preparation

#### DNA sample for FCS and smFRET

Fluorophore-labeled oligonucleotides, as described in ref.(*58*), were obtained from IBA (Göttingen, Germany). The DNA single strands were annealed using the following protocol: A 100 μL solution of two complementary single-stranded DNAs (ssDNA) at a concentration of 1 μM was heated to 95°C for 4 minutes and then cooled down to 4 °C at a rate of 1 °C/min in an annealing buffer (500 mM sodium chloride, 20 mM TRIS-HCl, and 1 mM EDTA at pH = 8).

#### Expression and purification of proteins

MalE single and double cysteine variants as well as the SBD2 (T369C) protein(*63*) were expressed and purified generally following established and published protocols(*55, 64, 65*) .For all MalE derivatives the T7/lac bacterial expression vector pET-23b(+) was chosen with a C-terminal His_6_-Tag. For SBD2 (T369C) an araBAD based bacterial expression system was selected including a N-terminal His_10_-Tag extension.

*E. coli* BL21(DE3)pLysS competent cells were transformed with the respective vector containing the DNA coding sequences for MalE and SBD2 variants. Expression cultures of the transformant cells were started in 2 l LB medium supplemented with carbenicillin (0.1 mg/ml), chloramphenicol (0.05 mg/ml), and 1 % D-glucose and grown at 37 °C until an optical density (OD_600_) of 0.6 – 0.8 was reached. Subsequently, over-expression was initiated by addition of 0.25 mM isopropyl β-D-1-thiogalactopyranoside (IPTG) for pET-23b(+) and using 0.2% L-arabinose for araBAD. The cells were harvested after 2 h and resuspended in 50 ml lysis buffer (50 mM Tris-HCl pH 8.0, 1 M KCl, 10 % glycerol, 10 mM imidazole) supplemented with 1 mM dithiothreitol (DTT). The collected cells were subjected to a 30-minute incubation at 4°C with 500 μg/ml DNase I, along with one tablet of EDTA-free protease inhibitor cocktail (cOmplete™, Roche) per 50 ml of culture extract. Additionally, 0.2 mM phenylmethylsulfonyl fluoride (PMSF) and 1 mM DTT were included. The cells were then lysed using an ultrasonic homogenizer (Digital Sonifier 250, Branson) equipped with a 5 mm diameter micro-tip probe, with parameters set at 25% amplitude, a total exposure time to ultrasound of 10 minutes, and time lapses of 0.5 seconds for ON/OFF pulse switches. Coarse cell debris were removed by centrifugation at 5000 × *g* for 30 min at 4 °C, followed by an ultracentrifugation step at 208,400 × *g* for 1 h at 4 °C to remove insoluble cellular components. The over-expressed proteins were purified by metal affinity chromatography using a Nickel functionalized agarose medium (Ni^2+^-Sepharose™ 6 Fast Flow, Cytiva), pre-equilibrated with 5 CV lysis buffer. A total of 50 ml cell lysate supernatant was loaded onto 4 ml of the Ni^2+^-Sepharose™ resin suspension and incubated overnight at 4 °C. Following the immobilization, the proteins were washed twice using two times 5 CV lysis buffer supplemented with 1 mM DTT and 0.2 mM PMSF and eluted with 1 CV elution buffer (50 mM Tris-HCl pH 8.0, 50 mM KCl, 10% glycerol, 250 mM imidazole, 0.2 mM PMSF and 1 mM DTT). To eliminate the excess of imidazole, the pooled protein containing fractions were each dialyzed overnight at 4°C against 100 volumes of dialysis buffer (50 mM Tris-HCl pH 8.0, 50 mM KCl) with the addition of 1 mM DTT, using a 10 kDa MWCO dialysis membrane tubing (SnakeSkin™, ThermoScientific). Next, the dialysis buffer was exchanged with 100 volumes of storage buffer (50 mM Tris-HCl pH 8.0, 50 mM KCl, 50% glycerol) supplemented with 1 mM DTT and the proteins were left to dialyze overnight at 4°C. The obtained proteins were snap frozen in liquid nitrogen and stored at -80°C until further use. The final concentrations were determined at a micro-volume Spectrophotometer (NanoPhotometer® N60, IMPLEN) using molar extinction coefficients of 66350 l*mol^-1^*cm^-1^ for MalE derivatives and 30370 l*mol^-1^*cm^-1^ for SBD2.

#### Labelling and labelled protein purification

The stochastic maleimide labelling and purification followed an already established protocol(*55, 64, 65*). For each labeling reaction 600 µg of protein from frozen stocks were used. His_6_-tagged MalE (S352C) and MalE (T36C-S352C) as well as the His_10_-tagged SBD2 (T369C) were incubated in labelling buffer (MalE variants: 50 mM Tris-HCl pH 7.4, 50 mM KCl; SBD2: 50 mM Tris-HCl pH 7.6, 150 mM NaCl) supplemented with 1 mM DTT to retain the reduced state of the introduced cysteine residues. In a first step, the proteins were immobilized by metal affinity on a Nickel functionalized agarose medium (Ni^2+^-Sepharose™ 6 Fast Flow, Cytiva), subsequently the maleimide reaction with 25 nmol of Cy3B for MalE (S352C) and SBD2 (T369C) (samples for FCS measurements) or with the combination of 25 nmol of each the Alexa Fluor 555 (ThermoFisher) and the Alexa Fluor 647 (ThermoFisher) for MalE (T36C-S352C; smFRET measurements) was carried out in the protein specific labelling buffer overnight at 4°C. The labeled, resin-bound proteins were washed with 1 CV of the respective labelling buffer and eluted with 500 µl labelling elution buffer (MalE variants: 50 mM Tris-HCl pH 8.0, 50 mM KCl, 500 mM imidazole; SBD2: 50 mM Tris-HCl pH 7.6, 150 mM NaCl, 500 mM imidazole). Following the maleimide labelling, the single, Cy3B labelled proteins were purified by size-exclusion chromatography (ÄKTA pure™ chromatography system, Cytiva; Superdex™ 75 Increase 10/300 GL, Cytiva), the MalE (T36C-S352C) was purified by anion exchange chromatography IEX (ÄKTA pure™ chromatography system, Cytiva; MonoQ™ 5/50 GL column, Cytiva).

Since the IEX purification will be topic of a forthcoming publication, we provide a short synopsis here. For MalE (T36C-S352C), the eluate from the maleimide labelling protocol was prepared for further purification by removal of the remaining KCl and imidazole from the labelling elution buffer that could otherwise interfere with the anion exchange process. This step was done using a Sephadex G-25 medium (PD MiniTrap™ G-25, Cytiva). The labelled protein was then eluted in 1 ml of anion exchange sample buffer (10 mM Tris-HCl pH 7.5). The anion exchange column was set up with a 5-CV H_2_O_dd_ wash, a 10-column volume equilibration with anion exchange sample buffer, 10-column volume equilibration with anion exchange elution buffer (10 mM Tris-HCl pH 7.5, 1 M NaCl) and a final 20-column volume equilibration with anion exchange sample buffer. The wash and all equilibrations were done with a 1 ml/ min flow rate. Labelled MalE (T36C-S352C) was loaded onto the column with 0.5 ml/ min, subsequently the resin-bound protein was washed with 10 column volumes anion exchange sample buffer and a 1 ml/ min flow rate. For the consecutive elution a linear increase in anion exchange elution buffer ratio with a slope corresponding to 7.5 mM NaCl per column volume was chosen. The flow rate was adjusted to 0.5 ml/ min. Fractions containing MalE (T36C-S352C) with both fluorophores (Alexa Fluor 555 and Alexa Fluor 647) in nearly stoichiometric amounts were selected and used for further analysis.

#### Sample immobilization (STORM)

For every experiment, an ibidi µ-Slide 8 Well Glass Bottom chamber was used. A single µ-Slide chamber was washed three times with 500 µl of PBS before each experiment. Then, 200 µl of a BSA-biotin solution (1 mg/mL in PBS (140 mM NaCl, 10 mM phosphate buffer, and 3 mM KCl, pH 7.4)) with 100 nm tracking fluorescent beads (TetraSpeck, Thermo Fisher) were added to the µ-Slide chamber and incubated for 10 minutes. The BSA-biotin was removed, and the chamber was carefully washed three times with 500 µl PBS. Next, a 200 µl streptavidin solution (1 mg/mL in PBS) was added to the µ-Slide and incubated for 10 minutes. The streptavidin solution was carefully removed, and the chamber was washed three times with imaging buffer IB (PBS with 10 mM magnesium chloride). A 1-10 µl solution of DNA origami nanorods (Gatta-STORM 94R, Gattaquant) was diluted with 200 µl IB and added to the µ-Slide chamber. The DNA origamis were incubated until a density of 1/µm^2^ was reached. Subsequently, the DNA origami solution was removed, and the chamber was washed three times with 500 µl IB. The photoswitching of the fluorophores during imaging was achieved by adding 500 µl of an oxygen scavenging system buffer(*63*) (pyranose oxidase at 3 U/mL, catalase at a final concentration of 90 U/mL, and 40 mM glucose in PBS) mixed with 0.1% (v/v) ß-mercaptoethanol.

### DNA-PAINT

DNA-PAINT samples were obtained ready to use by Gattaquant (Gatta-PAINT 80RG, Gattaquant, Germany).

#### 3D printing of the Brick-MIC platform

All models were designed and conceived using Onshape version 1.114 - 1.172. 3D printing was carried out using PLATech filament (OLYMPfila) on an Ultimaker +2 Extended fitted with a 0.4 mm nozzle. All models were printed with an infill density of 17%, three layers for outer walls and with a layer height of 0.1 mm. The printing speed was set to 50 mm/sec, and the nozzle temperature was maintained at 210 °C. To prevent warping, all parts were printed with a brim and without any supports. All models are available as STL file as Supplementary CAD files.

### Data Acquisition & Data analysis

#### µFCS

All samples were studied by positioning the confocal excitation volume into a 100 µL PBS droplet with concentrations ranging from 5 to 10 nM on a coverslip passivated with BSA (1 mg/ml in PBS).

The experimental setup shown in Figure 2 employed a 532 nm wavelength CW laser diode (5 mW output; CPS532, Thorlabs) as excitation light source. The beam underwent filtration through a clean-up filter (FL05532-10 ⌀12.5 mm, Thorlabs), attenuation via a continuous neutral density filter wheel (NDC-50C-2M, Thorlabs), and expansion using a telescope comprising a bi-concave lens (f = -50 mm, KBC043AR.14, Newport) and a plano-convex lens (f = 150 mm, LA1433-A-ML, Thorlabs). A dichroic beam splitter with high reflectivity at 532 nm (ZT532/640rpc, Chroma, USA) separated the excitation and emission beams to and from a high numerical aperture (NA) apo-chromatic objective (60X, NA 1.2, UPlanSAPO 60XW, Olympus, Japan). The emitted fluorescence was collected by the same objective, directed via a mirror into a piezo-directed optical mount (AG-M100N, Newport) through an inversely mounted 12 mm reflective collimator (RC12FC-P01, Thorlabs), which focused and coupled the emission beam into a multimode optical fiber (10 µm fiber core diameter, M64L01, Thorlabs). The fiber directed the emission light into a detection box, where it was collimated with a fixed-focus collimator (F220FC-532, Thorlabs) and then spectrally split into two separate photon streams by a dichroic mirror (ZT640rdc longpass, Chroma, USA). Individual photon streams were filtered with bandpass filters (for the green channel: FF01-582/75-25 ⌀25 mm, Semrock Rochester NY, USA; for the red channel: ET700/75m, Chroma) and detected by two distinct photomultiplier tubes with different spectral sensitivities (for the green channel: H10682-210, Hamamatsu, Japan; for the red channel: H10682-01, Hamamatsu, Japan). The detector outputs for FCS analysis were either recorded by a NI-Card PCI-6602 (National Instruments, USA) using LabView data acquisition software from the Weiss laboratory(*66*), or via a counter/timer device module (USB-CTR04, Measurement Computing, USA) with custom-made acquisition software written in Python, available for download as compiled executable or editable python code at https://github.com/harripd/mcc-daq-acquisition. We provide information on which counter module was used for measurements in the respective text and figure caption.

Data analysis was performed using a home-written Python script(*41*) (https://github.com/PSBlmu/FCS---analysis), where the intensity fluctuation of freely diffusing molecules *F*(τ) is analyzed via autocorrelation:

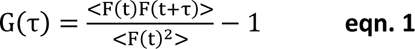

Here, the correlation amplitude *G*(*τ*) describes the self-similarity of the signal in time. Average fluorescence intensities at time points t and later lag times *τ* are used for analysis. *G*(*τ*) is analyzed with a 3D diffusion model:

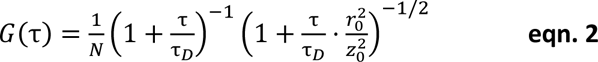

Here, *N* represents the average number of molecules in the confocal volume, τ_*D*_ is the average diffusion time, and *r*_0_, *z*_0_ define the lateral and axial radial distances of the detection volume, which define the geometry parameter 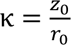

#### µALEX

All samples for smFRET analysis were measured in a 100 µL PBS droplet with concentrations ranging from 50 to 100 pM on a coverslip passivated with BSA (1 mg/ml in PBS). The experimental setup utilized alternating laser excitation (ALEX) of two diode lasers: OBIS 532-100-LS (Coherent, USA), operated at 60 µW for donor molecules at 532 nm, and OBIS 640-100-LX (Coherent, USA), operated at 25 µW for acceptor molecules at 640 nm, both in alternation mode with a 100 µs period. The lasers were combined by an aspheric fiber port (PAF2S-11A, Thorlabs, USA), coupled into a polarization-maintaining single-mode fiber P3-57 488PM-FC-2 (Thorlabs, USA) and collimated (RC12APC-P01, Thorlabs, USA) before entering an epi-illuminated confocal microscope (Olympus IX71, Hamburg, Germany). Excitation and emission collection was done by the same water immersion objective (60X, NA 1.2, UPlanSAPO 60XW, Olympus, Japan) and spectral separation was achieved by a dual-edge beamsplitter ZT532/640rpc (Chroma/AHF, Germany). Fluorescence emitted from the sample was collected by the same objective and further focused via an achromatic lens (AC254-200-A, Thorlabs) directly into the single-photon avalanche diodes (PDM 50-Micron, MPD). The small active area of the detectors (ø 50 µm) served as a pinhole. Before that, the photon streams were spectrally split into the donor and acceptor channels by a single-edge dichroic mirror H643 LPXR (AHF, Germany). Fluorescence emission was filtered by band-pass filters directly in front of each detector: for the donor channel FF01-582/75-25 (Semrock/AHF, Germany) and ET700/75m Chroma (AHF, Germany) for the acceptor channel. The detector outputs for µsALEX were recorded by an NI-Card PCI-6602 (National Instruments, USA) using LabView data acquisition software from the Weiss laboratory(*66*).

Data analysis was performed using a home-written software package, as described previously(*65*). Single bursts were identified first using All-Photon-Burst-Search (APBS) with a threshold for burst start/stop of 15 photons(*40*), a time window of 500 µs, and a minimum total photon number of 150 within the burst. Based on these data donor leakage (α) and direct acceptor excitation (δ) were determined as mean values from a 1D-fit of background-corrected donor-only E* and acceptor-only S* distributions. Then, a Dual-Channel-Burst-Search (DCBS) was performed with similar parameters to determine excitation flux (β) and detection efficiency and quantum yields (γ)(*42*). E-histograms of double-labelled FRET species were generally extracted by selecting 0.3 < S < 0.7. E-histograms were fitted with a Gaussian function according to 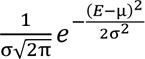, where *E* represents the measured FRET efficiency for every detected molecule, μ is the mean, and σ the standard deviation. For FRET efficiency to inter-dye distance conversion, the Förster equation was used:

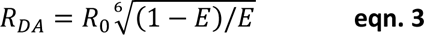

The Förster radius *R*_0_ is given by the following equation(*67*):

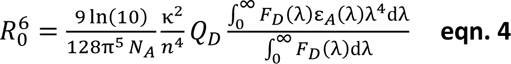

where *N_A_* is the Avogadro constant, *κ*^2^ the dipole orientation factor, *n* the averaged refractive index of the medium, *Q_D_* the donor quantum yield, *F_D_* the donor emission spectrum, and the acceptor absorbance spectrum *ε_A_*. We used values of R_0_ known from the literature: Cy3-ATTO647N: R_0_ = 5.1 nm(*56, 57*); Cy3B-ATTO647N: R_0_ = 6.7 nm(*57–59*). For Cy3B-Cy5 we used an unpublished approach to determine R_0_ = 7.4 nm by least-square fitting of the data (E values) to the known inter-dye distances of donor and acceptor on the dsDNA(*58*) using equation 3.

#### TCSPC

The setup for fluorescence lifetime decay measurements combines the excitation layer of the ALEX modality with the emission layer of the FCS modality (see above, Supplementary Video 3). The configuration utilized a 532 nm wavelength fiber-coupled pulsed laser operating at a rate of 20 MHz and a power of 55 µW (LDH-P-FA-530B with PDL 828 ’Sepia II’ controller, Picoquant, Germany), detected by PMTs connected to a Multiharp 150 8N (Picoquant, Germany). All samples were measured in a 50 µL PBS droplet with concentrations of 10 nM on a coverslip. The Cy3B and Alexa 546 fluorophore-labeled dsDNA samples were measured for 10 minutes, and the Atto 550 sample was measured for 60 minutes. The IRF was acquired by measuring a 50 µL PBS droplet for 60 minutes. Lifetime decays were determined using the SymPhoTime 64 analysis program and fitted using a single Exponential Tailfit model (n = 1). For the Cy3B and Alexa 546 samples, a threshold window of 21.79 ns was used for the fitting, and for the Atto 550 samples, a window of 16.15 ns was used (see Supplementary Figure 6).

#### µEpi

The setup for widefield imaging utilizes an external fiber-coupled laser box (READY BeamTM ind 2 1007773, Fisba, Switzerland). A 640 nm wavelength continuous-wave excitation laser was used which provides an output of 30 mW (measured after the objective). The beam was collimated through a parabolic mirror (RC04APC-P01, Thorlabs), passes through a clean-up filter (ZET 635/10 ⌀25 mm, Chroma), and was expanded with a plano-concave lens (f = -25 mm, LC1054-ML - Ø1/2, Thorlabs). A plano-convex lens (f = 60 mm, LA1134-A-ML - Ø1, Thorlabs) focuses the beam into the back focal plane of the objective. A dichroic beam splitter with high reflectivity at 640 nm (ZT532/640rpc, Chroma, USA) separates excitation and emission beams into and from a high numerical aperture (NA) apo-chromatic oil immersion objective (60X, NA 1.35, UPlanSApo60XO, Olympus, Japan). Fluorescence emitted from the sample was collected by the same objective and was further focused via an achromatic lens (AC254-150-A-ML, Thorlabs), projecting a real image onto the chip of a CMOS camera (U3-30C0CP-M-GL rev.2.2, IDS)(*68*). The photon stream is further filtered with a band-pass filter (ET700/75m Chroma, AHF, Germany) before reaching the camera sensor.

The photoswitching of the fluorophores was achieved by adding 500 µl of an oxygen scavenging system buffer(*63*) (pyranose oxidase at 3 U/mL, catalase at a final concentration of 90 U/mL, and 40 mM glucose in PBS) mixed with 0.1% (v/v) ß-mercaptoethanol. The laser power used for imaging was 30 mW, which corresponds to approximately 37 kW/cm^2^ with an illuminated area of 80 x 80 µm^2^ (Supplementary Figure 17). Diffraction limited recordings were acquired using the original software that came with the cameras. All recordings were done at 10 frames per second, with an exposure time of 100 ms and the analog gain set at maximum.

Super-resolution image reconstruction was performed using the ImageJ plug-in Thunderstorm(*61*). Each localization was filtered using a B-Spline wavelet filter as described. The local maxima method was employed for the approximate localization of the molecules. Sub-pixel localization was achieved by fitting an Integrated Gaussian model using a weighted least squares method with a 3-pixel fitting radius. The super-resolution image was rendered with a pixel size of 12 x 12 nm^2^. For STORM, drift correction was implemented using the fiducial marker algorithm with fluorescent tetraspeck beads (100 nm, Thermo Fisher) on the sample and lateral drift was assessed using tracking of bead positions over ∼30 mins (Supplementary Fig. 18). The maximal search tracking distance was set at 20 nm, and the minimum visibility ratio was maintained at 0.9 per frame. For PAINT, drift correction was performed using a cross-correlation algorithm, correlating the positioning of all blinking events of each molecule in the movie over time.

## Acknowledgments

This work was financed by the European Commission (ERC-STG 638536 – SM-IMPORT to T.C.), the Bundesministerium für Bildung und Forschung (KMU grant „quantumFRET“ to T.C.), the Israel Science Foundation (grants 556/22 and 3565/20 to E.L.), NIH (grant R01 GM130942 to E.L. as subaward) and the Center for Nanosicence (CeNS). The authors also thank the Graduate School Life Science Munich (LSM) for support. N.Z. acknowledges a postdoctoral fellowship from the Alexander von Humboldt foundation. We thank C. Gebhardt and K. Schütze for programming support, M. Isselstein for valuable discussions, and J. Schneider and J. Piedra for experimental support. We thank J. Schmied and GATTAquant for the kind gift of the STORM nanoruler.

## Author contributions statement

G.G.M. and T.C. designed and conceived the study. G.G.M. conceived and designed the modular platform architecture. G.G.M., O.B., N.Z. and T.C. planned the layout of the microscopes. G.G.M., O.B. and N.Z. built microscopes. N.D.W. provided samples. P.K., P.D.H. and N.Z. wrote data acquisition and data analysis software. G.G.M., O.B. and J.R.L.P. conducted experiments and analyzed data. G.G.M. prepared figures. E.L. and T.C. acquired funding. T.C. supervised the study. G.G.M. and T.C. wrote the manuscript, which was reviewed, edited and approved by all authors.

## Data and Materials Availability

All data needed to evaluate the conclusions in the paper are present in the paper and/or the Supplementary Materials. Supplementary files including videos and 3D-printing templates are available in a repository under https://zenodo.org/records/10441063.

## Competing interest statement

G.G.M., O.B., N.Z. and T.C. have submitted a patent for commercialization of the Brick-MIC microscopy platform. G.G.M. and T.C. declare commercial interest in Brick-MIC. The authors declare no other competing interests.

## Supplementary Information

**Supplementary Figure 1:**
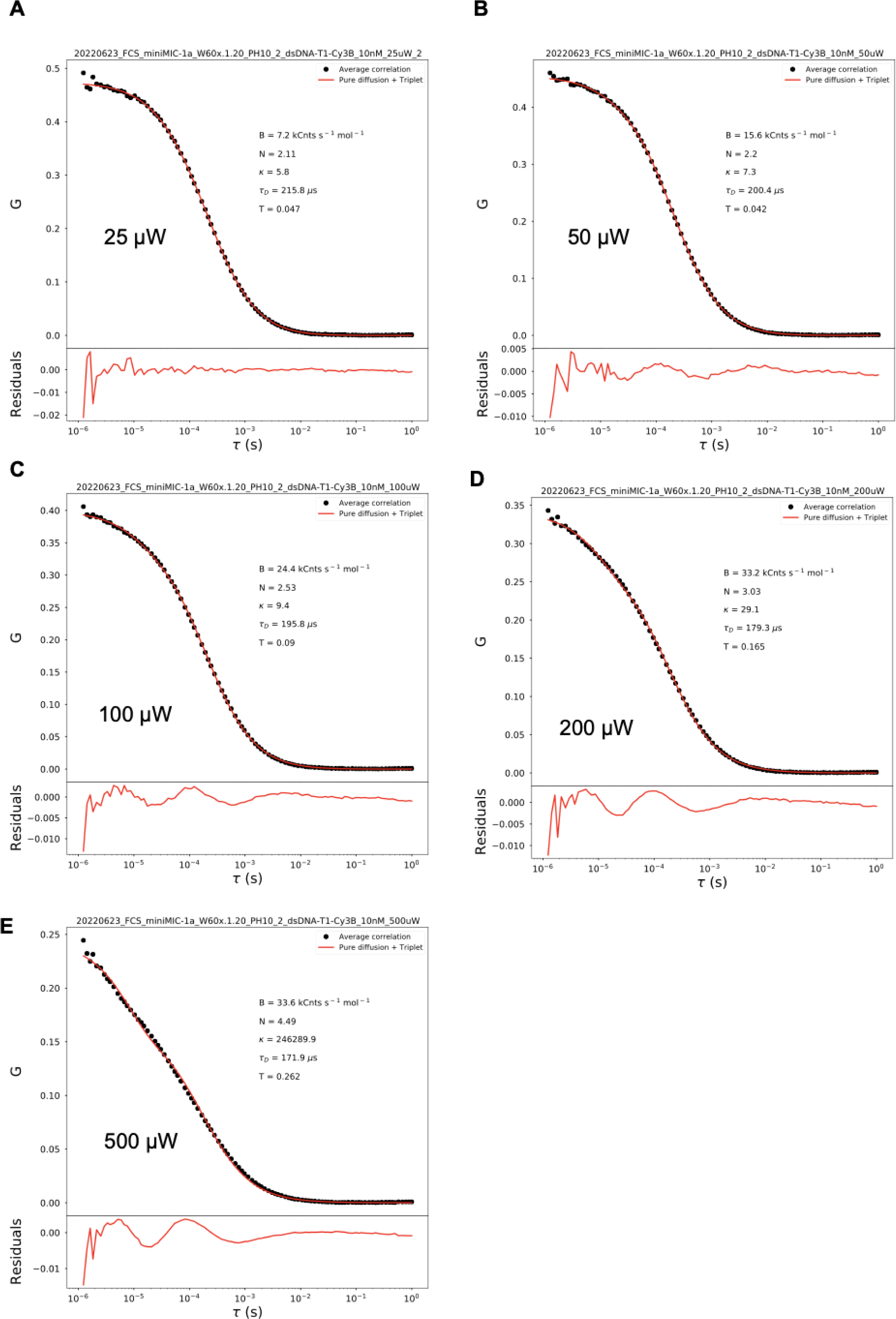
FCS laser power dependency of a 10 nM solution of a 40mer dsDNA sample labelled with Cy3B excited at 532 nm with the PMT variant by using a 10 µm pinhole.

**Supplementary Figure 2:**
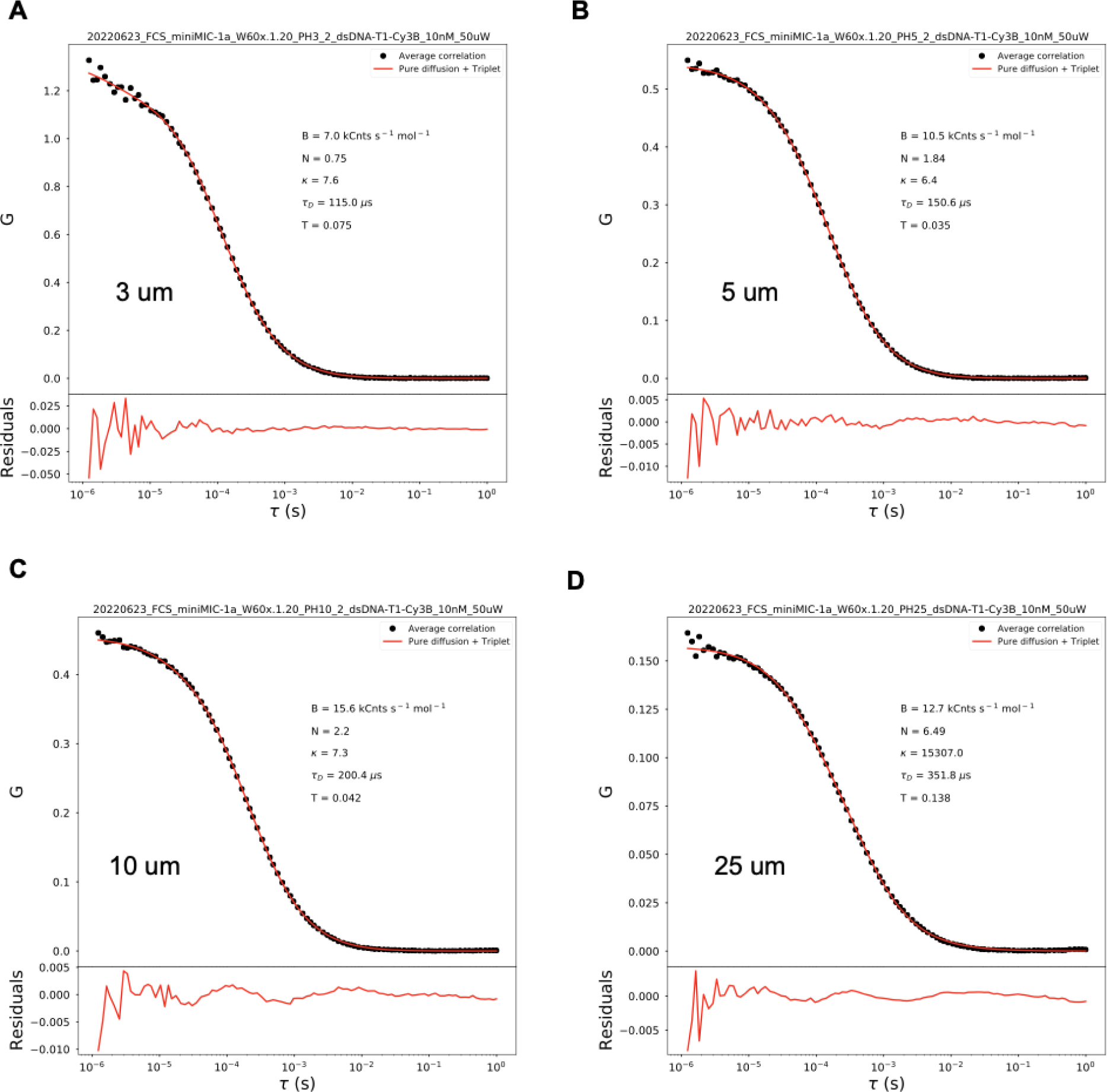
FCS pinhole size dependency of the PMT variant of a 10 nM solution of a 40mer dsDNA sample labelled with Cy3B and excited with 50 µW laser power at 532 nm.

**Supplementary Figure 3:**
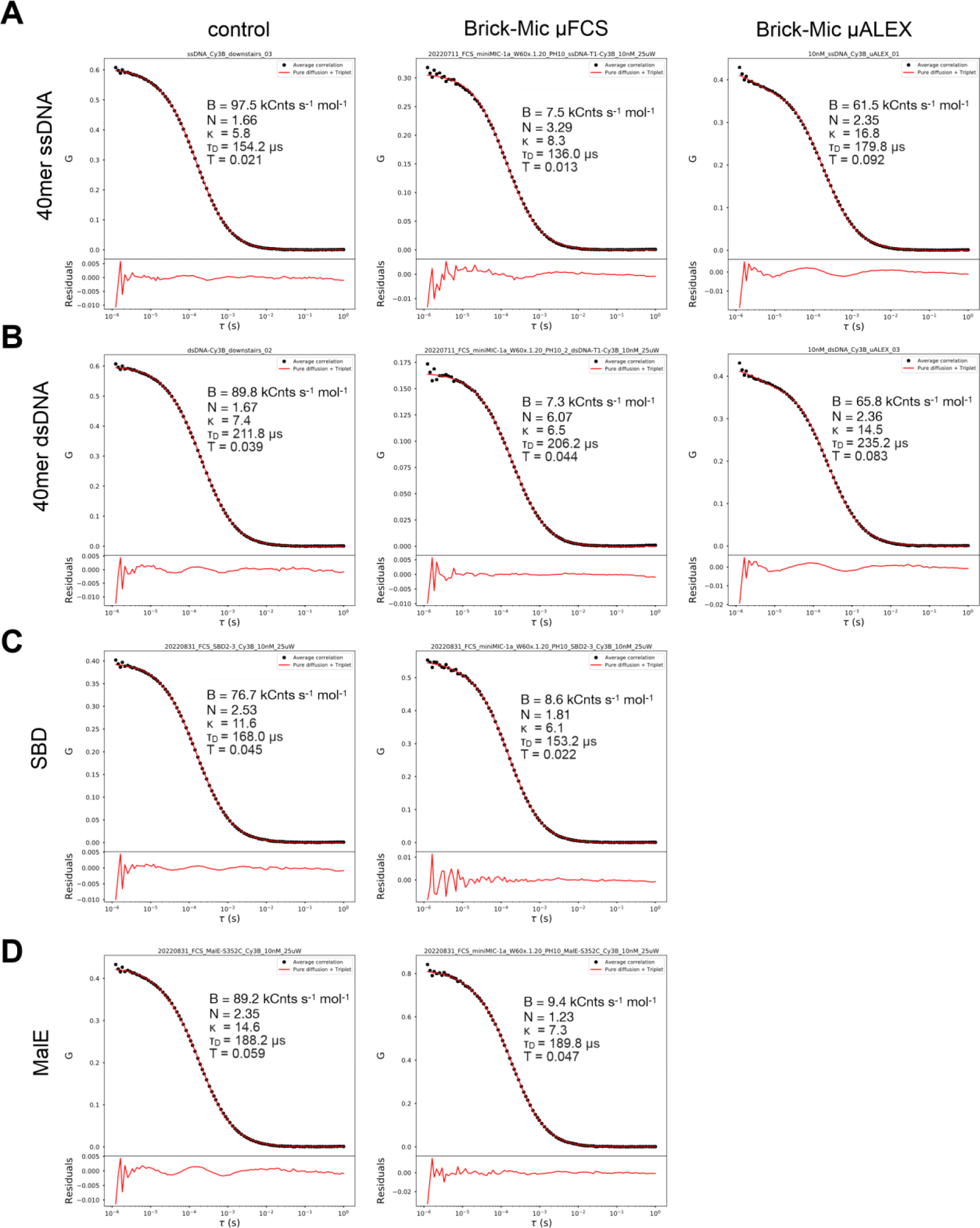
FCS comparison between a home-built confocal setup (left side) and two modalities of Brick-MIC: PMT variant using a 10 μm pinhole diameter (middle) and APD variant (right). A-D) show experiments and FCS curves of various biomolecules with different masses and hydrodynamic radii. All measurements were conducted at a 25 μW laser power with excitation at 640 nm.

**Supplementary Figure 4:**
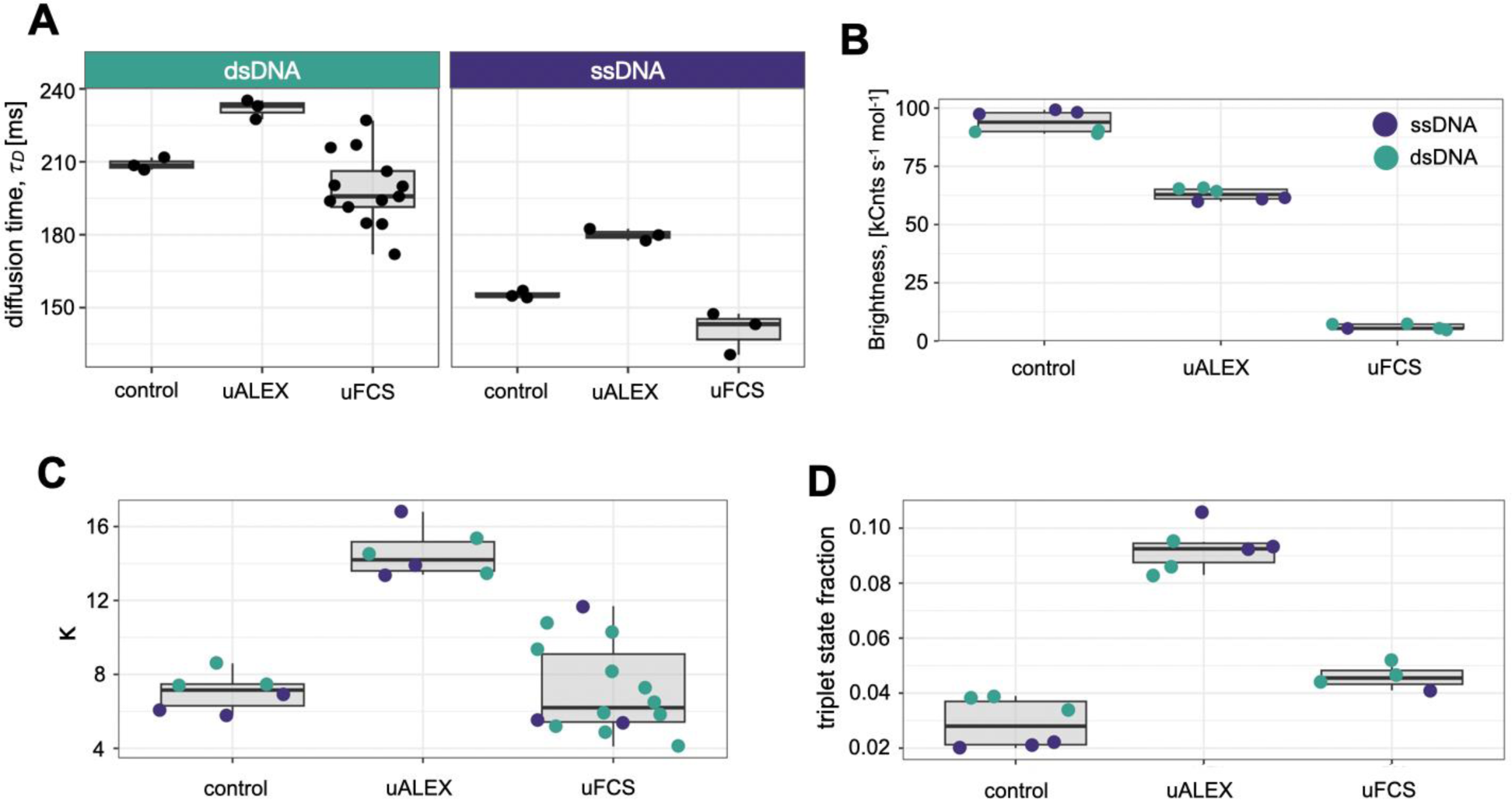
Boxplots of FCS fit values comparison shown in Supplementary Figure 3. Comparison between the a home-built confocal setup (control) and two modalities of Brick-MIC: the PMT variant using a 10 μm pinhole diameter (μFCS), and the APD variant (μALEX). A) diffusion times of dsDNA and ssDNA shown for all setups. B) molecular brightness, C) confocal structure parameter κ, and D) triplet state fraction.

**Supplementary Figure 5.**
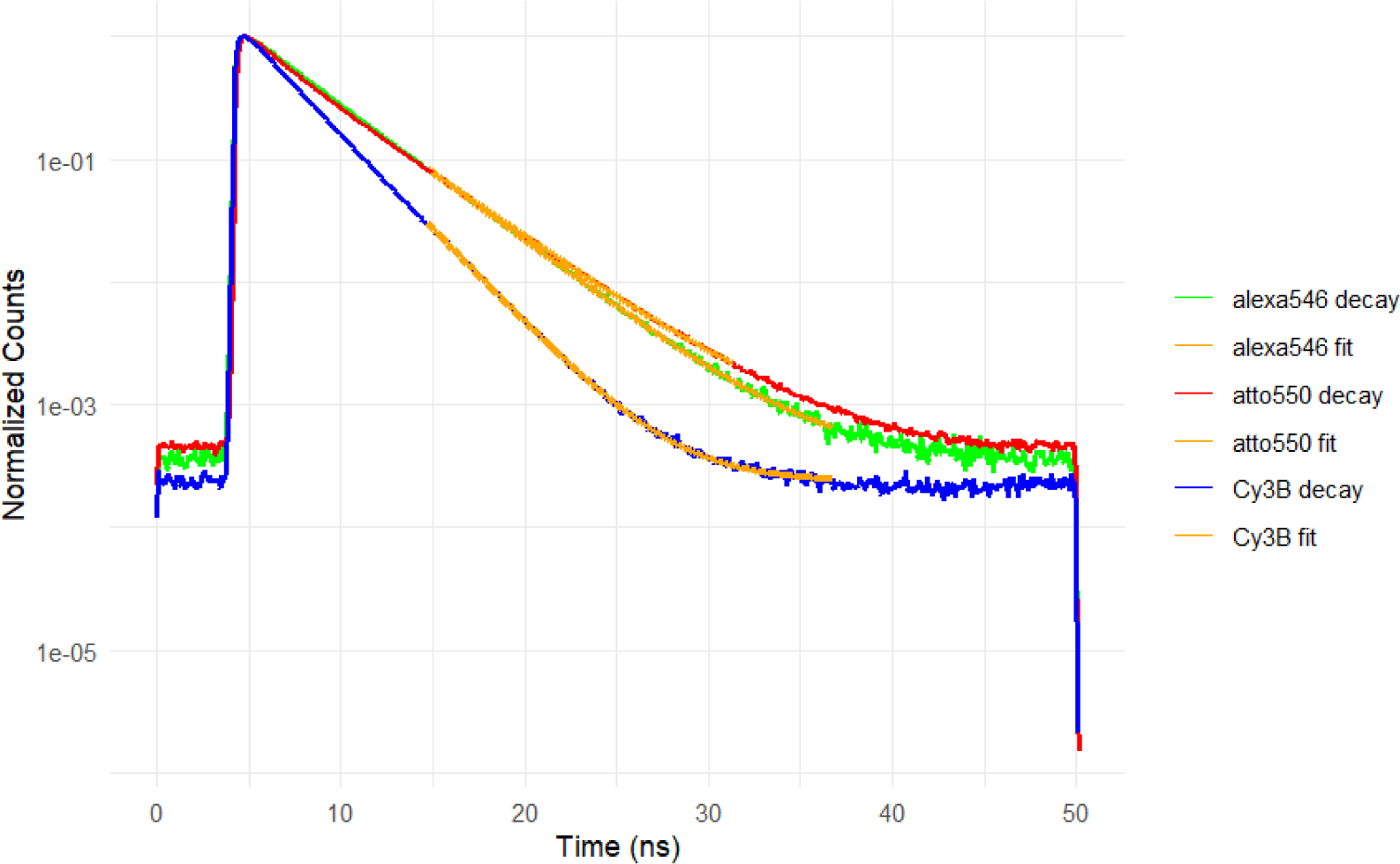
Fluorescence lifetime decays of fluorophore-labeled dsDNA. A) Normalized lifetime data and single-exponential fit of three fluorophores on dsDNA using the PMT version of the Brick-MIC.

**Supplementary Figure 6.**
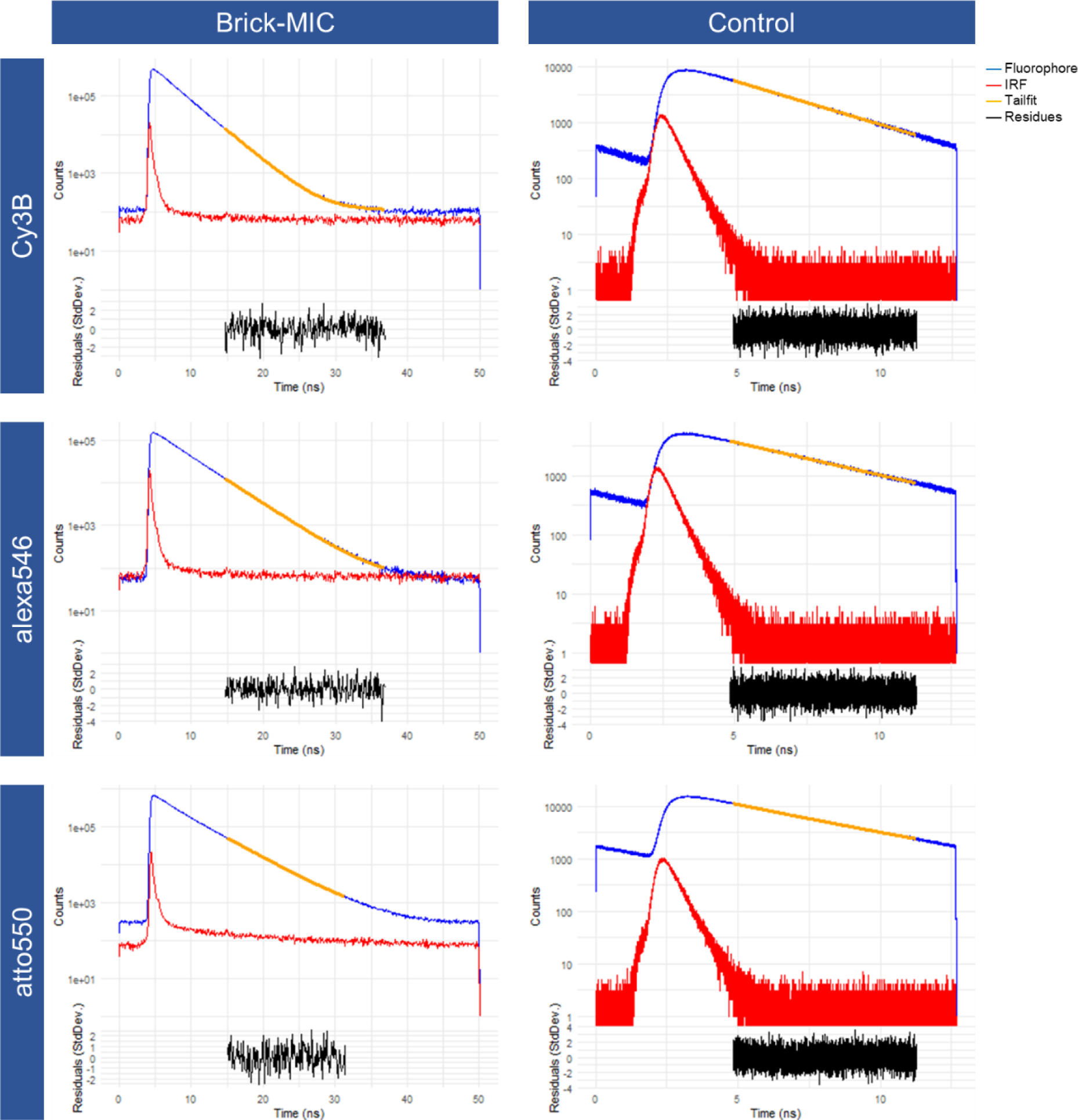
Lifetime measurements, exponential tailfit and residues of fluorophore-labeled dsDNA. TCSPC measurements of three dyes on dsDNA (rows) obtained with the Brick-MIC platform (left column) and a home built TCSPC setup^39^ (right column). The IRF FWHM where 0.32 ns and 0.65 ns for the Brick-MIC and the control, respectively.

**Supplementary Figure 7.**
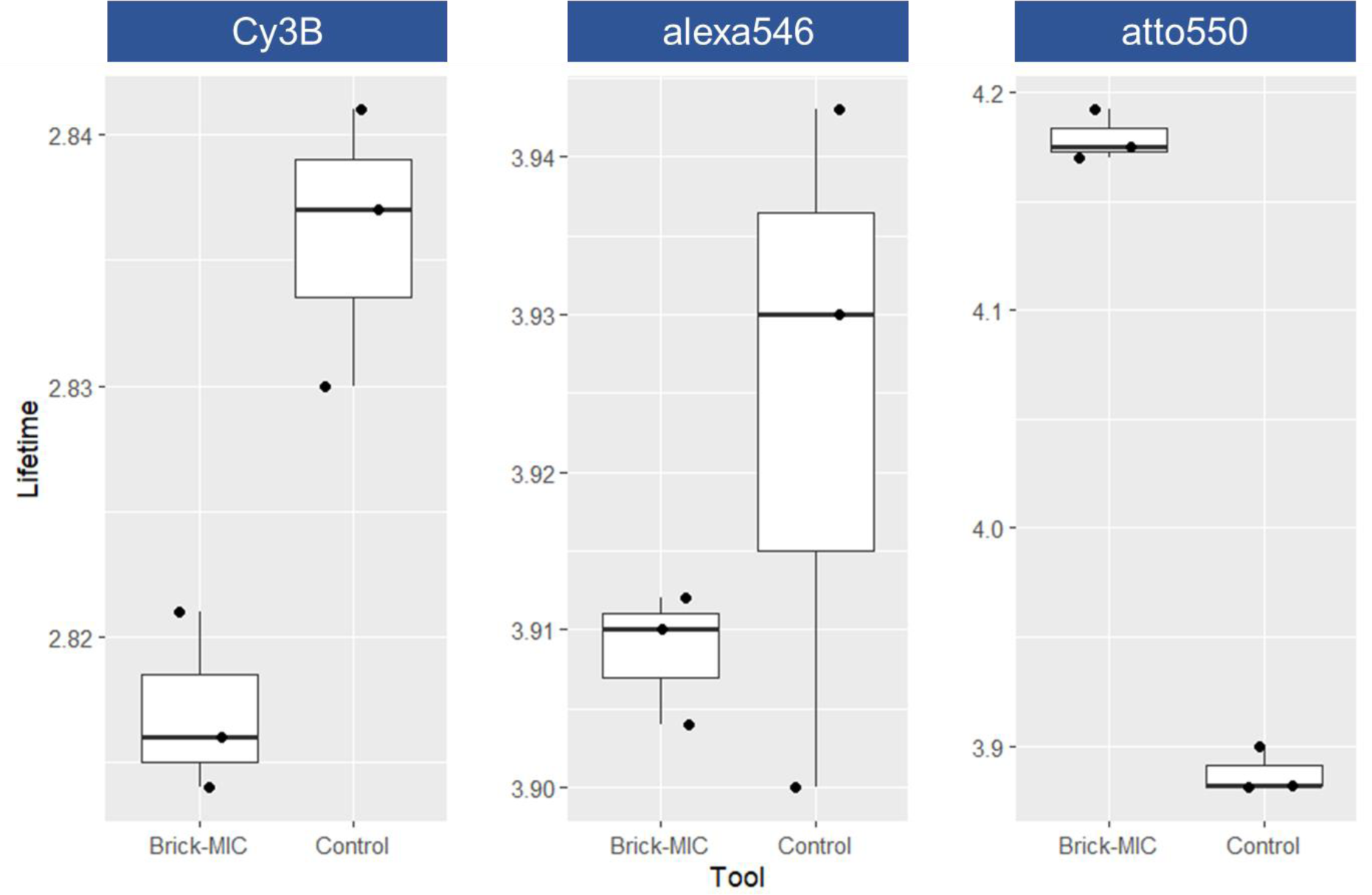
Boxplots of fluorescence lifetimes from three fluorophore-labeled dsDNA taken with the Brick-MIC platform and with a home built TCSPC setup^39^ (control).

**Supplementary Figure 8:**
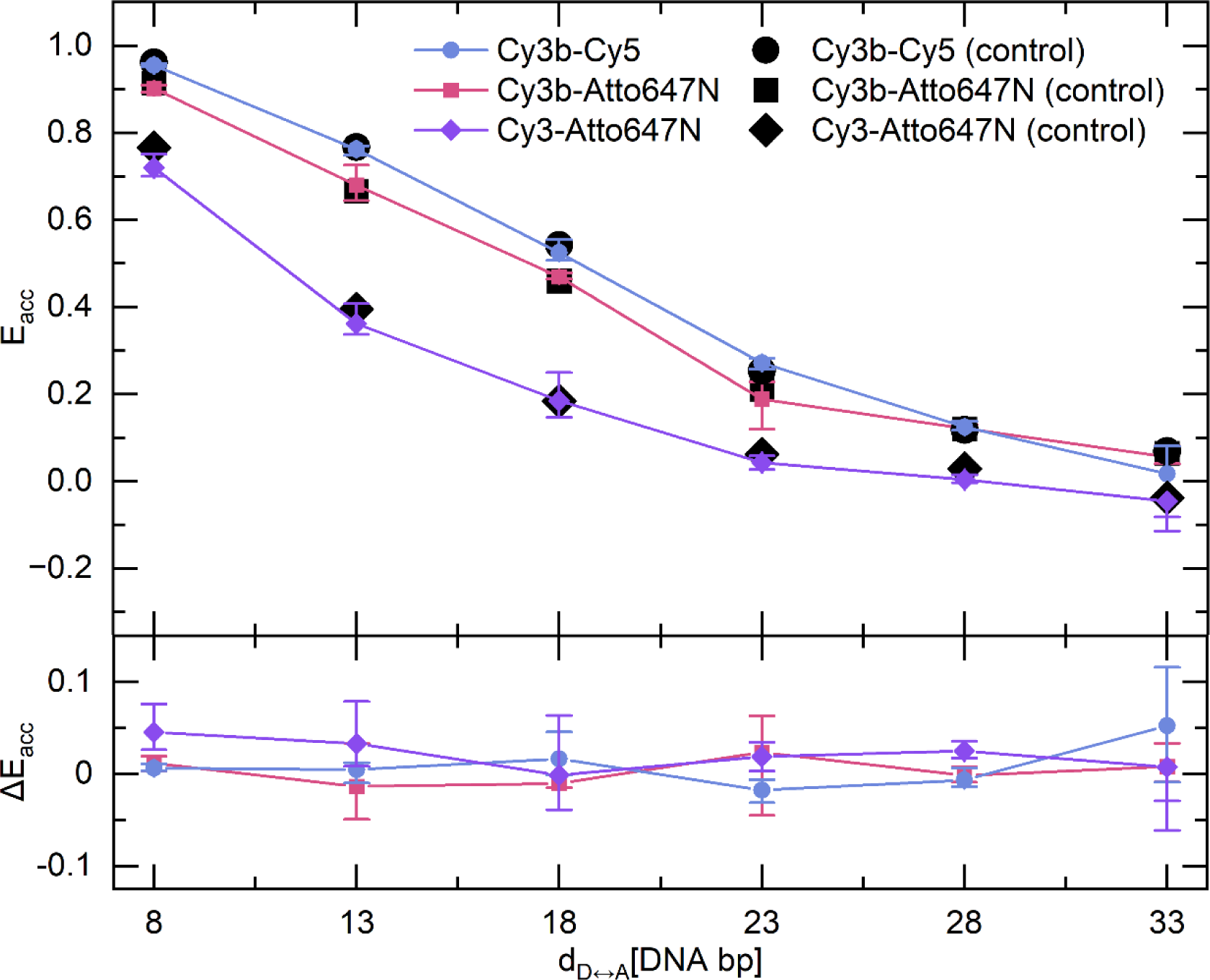
Accurate FRET values comparison between the Brick-MIC APD variant and a home-built confocal microscope. smFRET measurements were performed on donor and acceptor fluorophore pairs Cy3B-Cy5, C3b-Atto647N and Cy3-Atto647n which were covalently labelled to dsDNA (with distances between donor and acceptor fluorophore pairs in terms of DNA base pairs 8, 13, 18, 23, 28 and 33). Setup independent accurate FRET values were compared between the Brick-MIC ALEX modality (coloured symbols) and a home-built confocal microscope (black filled symbols).

**Supplementary Figure 9:**
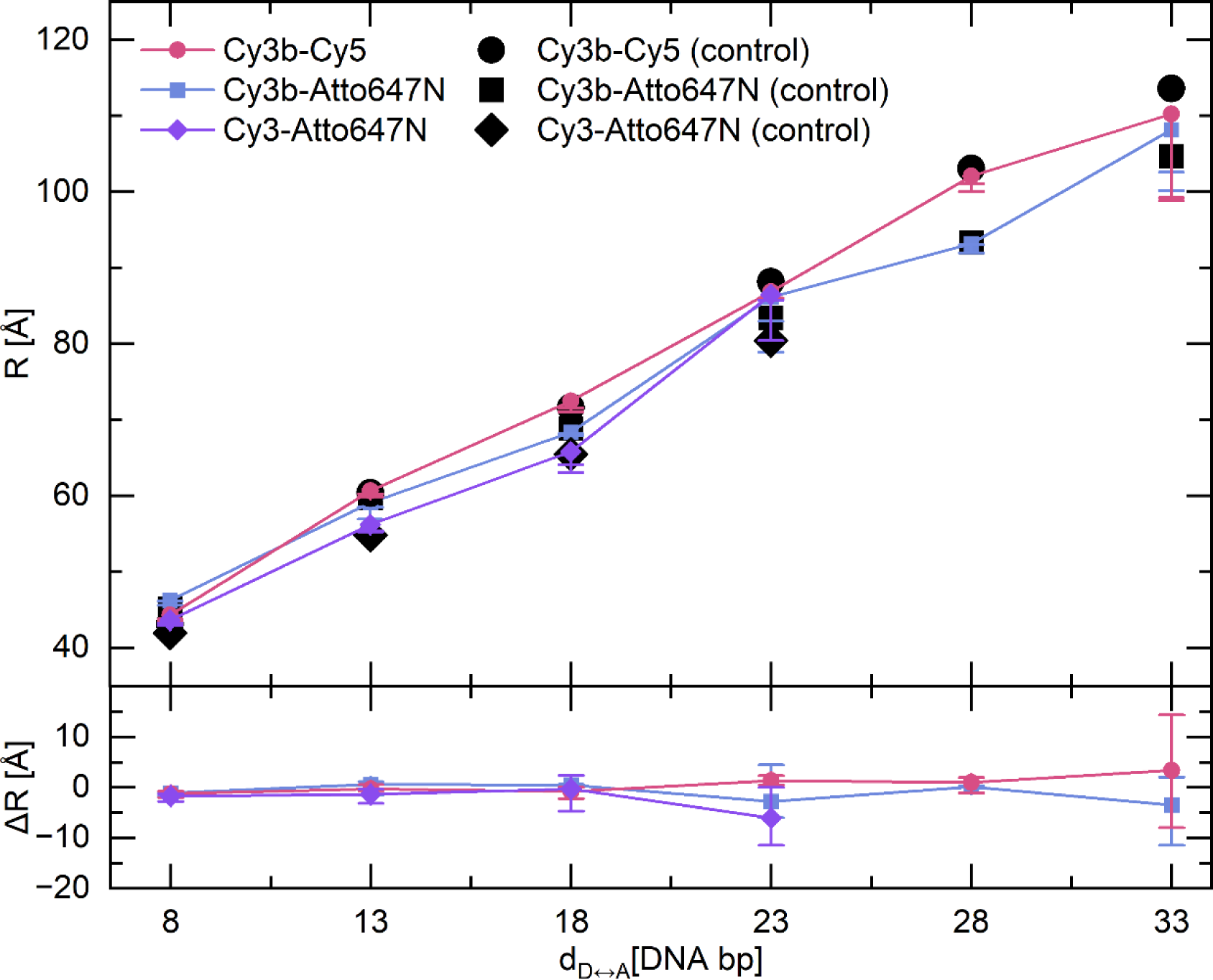
Calculated distances comparison between the Brick-MIC APD variant and a home-built confocal microscope. The setup independent accurate FRET values were used to calculate the distances in Å between donor and acceptor fluorophore pairs (i. e. Cy3B-Cy5, Cy3B-Atto647N and Cy3-Atto647N) subjected to the DNA base pair distances 8, 13, 18, 23, 28 and 33. Filled coloured symbols display smFRET measurements performed on the Brick-MIC and black filled symbols display measurements on a home-built confocal microscope .

**Supplementary Table 1:** Summary of fit results (2D Gaussian model) of smFRET data obtained with the Brick-MIC APD variant for a Cy3B and Atto647N dsDNA ladder.

| bp | 8 | 13 | 18 | 23 | 28 | 33 |
| --- | --- | --- | --- | --- | --- | --- |
| $E^*$ | 0.84 | 0.58 | 0.35 | 0.20 | 0.15 | 0.12 |
| $\sigma_{E^*}$ | 0.06 | 0.09 | 0.07 | 0.05 | 0.04 | 0.04 |
| $S^*$ | 0.54 | 0.57 | 0.64 | 0.66 | 0.69 | 0.69 |
| $\sigma_{S^*}$ | 0.07 | 0.07 | 0.07 | 0.06 | 0.06 | 0.06 |

**Supplementary Figure 10:**
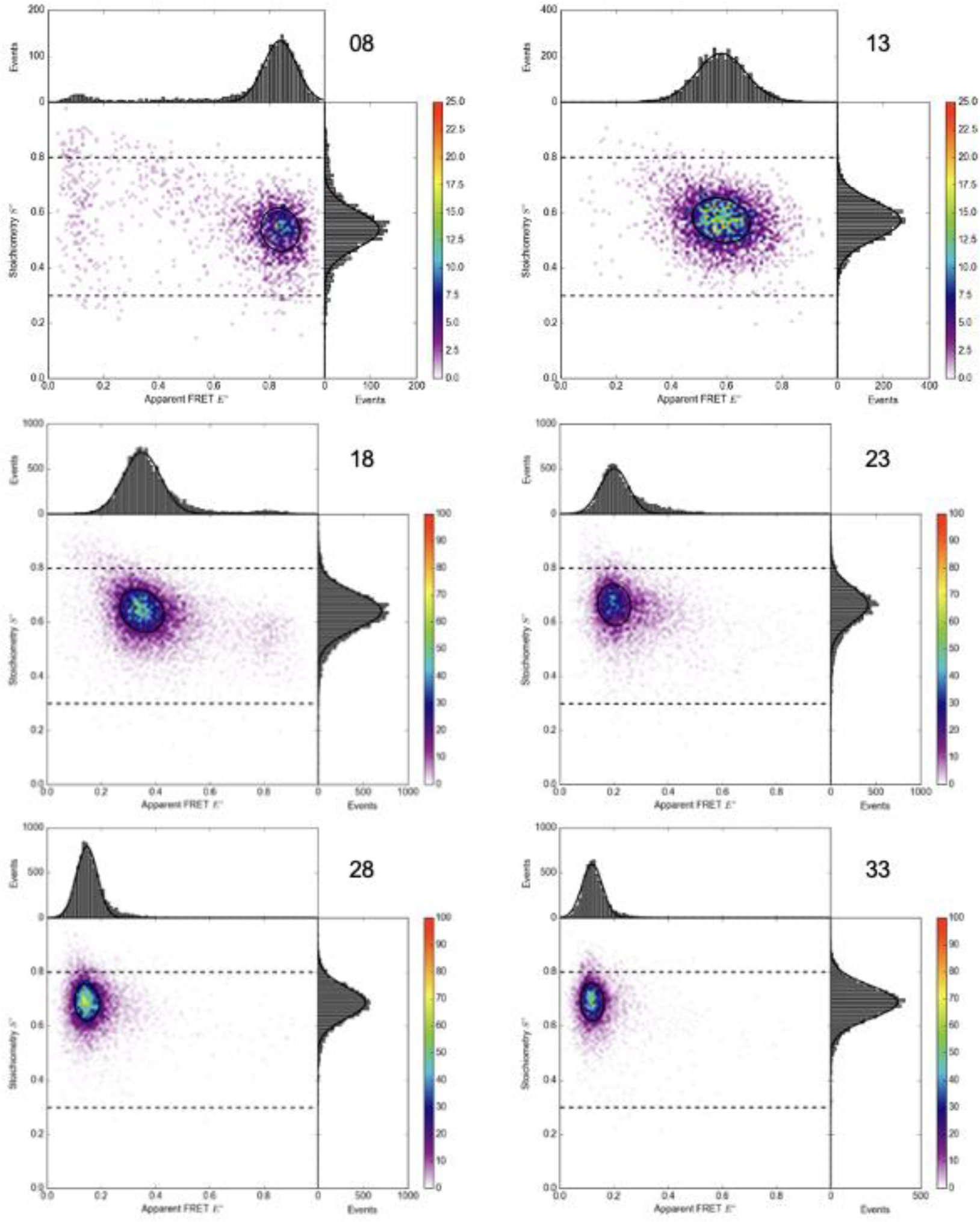
Two-dimensional ALEX-histograms showing setup-dependent uncorrected apparent E* vs S* values from single-molecule observation of dsDNA samples using the APD variant of Brick-MIC. The dsDNA ladder was labeled with Cy3B and Atto647N. The corresponding distances are 8 bp (top-left), 13 bp (top-right), 18 bp (middle-left), 23 bp (middle-right), 28 bp (lower-left) and 33 bp (lower-right).

**Supplementary Table 2:** Summary of fit results (2D Gaussian model) of smFRET data obtained with the Brick-MIC APD variant for a Cy3B and Cy5 dsDNA ladder.

| bp | 8 | 13 | 18 | 23 | 28 | 33 |
| --- | --- | --- | --- | --- | --- | --- |
| $E^*$ | 0.87 | 0.56 | 0.33 | 0.19 | 0.13 | 0.13 |
| $\sigma_{E^*}$ | 0.06 | 0.10 | 0.07 | 0.05 | 0.04 | 0.03 |
| $S^*$ | 0.59 | 0.64 | 0.73 | 0.76 | 0.81 | 0.79 |
| $\sigma_{S^*}$ | 0.06 | 0.07 | 0.07 | 0.06 | 0.05 | 0.05 |

**Supplementary Figure 11:**
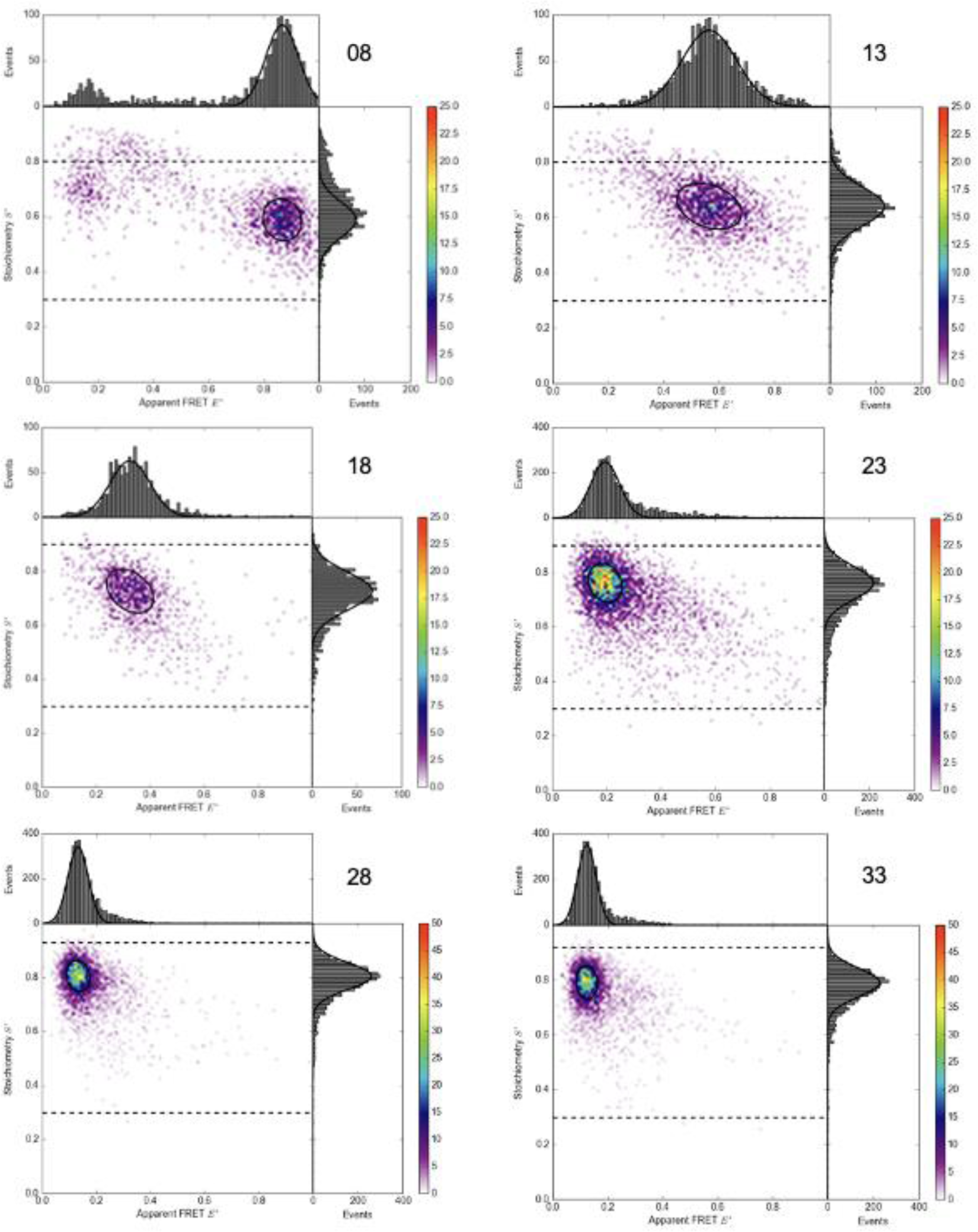
Two-dimensional ALEX-histograms showing setup-dependent uncorrected apparent E* vs S* values from single-molecule observation of dsDNA samples using the APD variant of Brick-MIC. The dsDNA ladder was labeled with Cy3B and Cy5. The corresponding distances are 8 bp (top-left), 13 bp (top-right), 18 bp (middle-left), 23 bp (middle-right), 28 bp (lower-left) and 33 bp (lower-right).

**Supplementary Table 3:** Summary of fit results (2D Gaussian model) of smFRET data obtained with the Brick-MIC APD variant for a Cy3 and Atto647N dsDNA ladder.

| bp | 8 | 13 | 18 | 23 | 28 | 33 |
| --- | --- | --- | --- | --- | --- | --- |
| $E^*$ | 0.84 | 0.58 | 0.36 | 0.21 | 0.17 | 0.14 |
| $\sigma_{E^*}$ | 0.06 | 0.08 | 0.08 | 0.07 | 0.06 | 0.05 |
| $S^*$ | 0.50 | 0.44 | 0.42 | 0.42 | 0.41 | 0.40 |
| $\sigma_{S^*}$ | 0.06 | 0.06 | 0.06 | 0.06 | 0.07 | 0.07 |

**Supplementary Figure 12:**
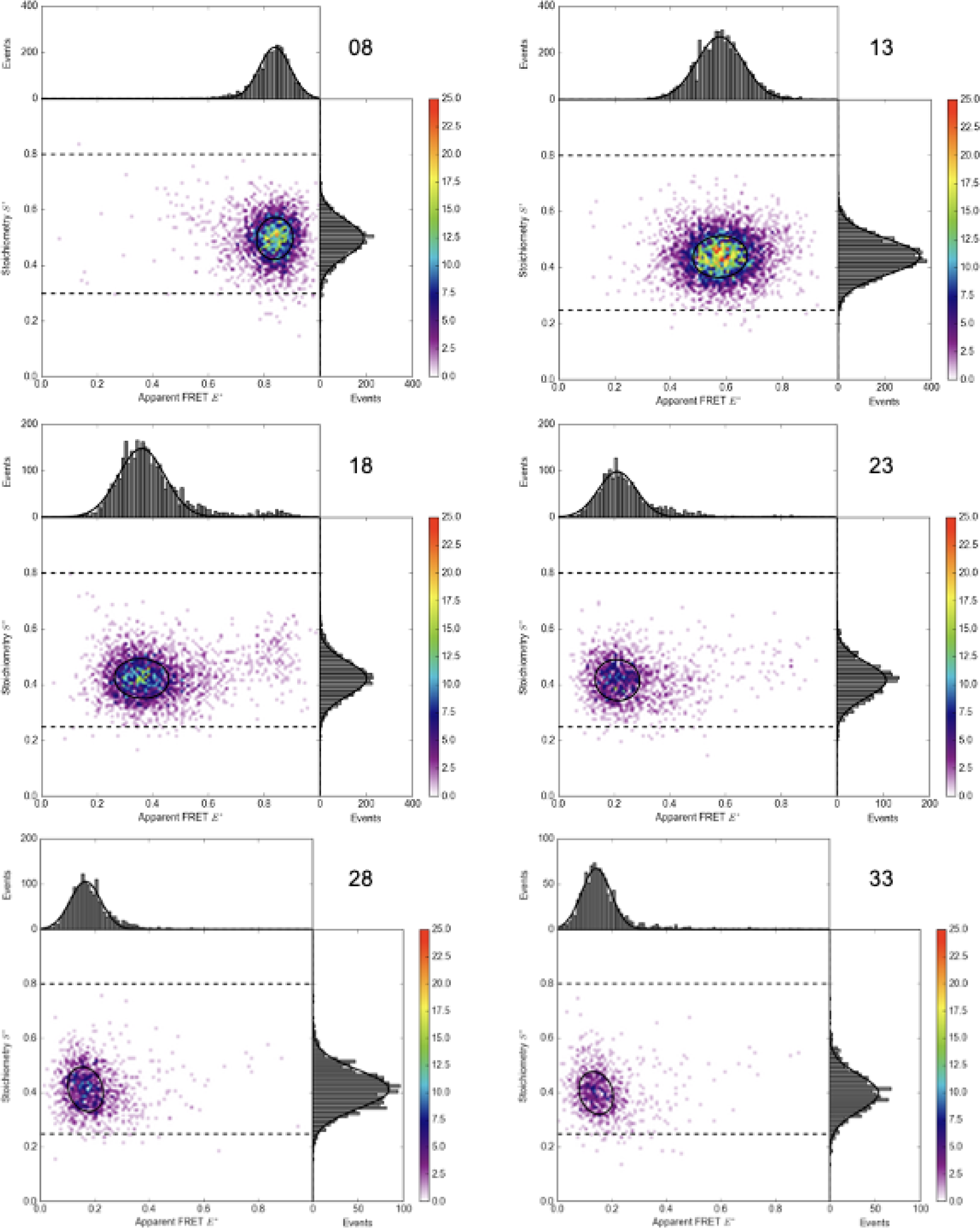
Two-dimensional ALEX-histograms showing setup-dependent uncorrected apparent E* vs S* values from single-molecule observation of dsDNA samples using the APD variant of Brick-MIC. The dsDNA ladder was labeled with Cy3 and Atto647N. The corresponding distances are 8 bp (top-left), 13 bp (top-right), 18 bp (middle-left), 23 bp (middle-right), 28 bp (lower-left) and 33 bp (lower-right).

**Supplementary Table 4:** Summary of fit results (2D Gaussian model) of smFRET data obtained with the home-built confocal microscopy for a Cy3B and Atto647N dsDNA ladder.

| bp | 8 | 13 | 18 | 23 | 28 | 33 |
| --- | --- | --- | --- | --- | --- | --- |
| $E^*$ | 0.90 | 0.65 | 0.47 | 0.28 | 0.22 | 0.19 |
| $\sigma_{E^*}$ | 0.04 | 0.08 | 0.07 | 0.05 | 0.04 | 0.04 |
| $S^*$ | 0.53 | 0.55 | 0.58 | 0.58 | 0.59 | 0.59 |
| $\sigma_{S^*}$ | 0.06 | 0.07 | 0.08 | 0.07 | 0.07 | 0.07 |

**Supplementary Figure 13:**
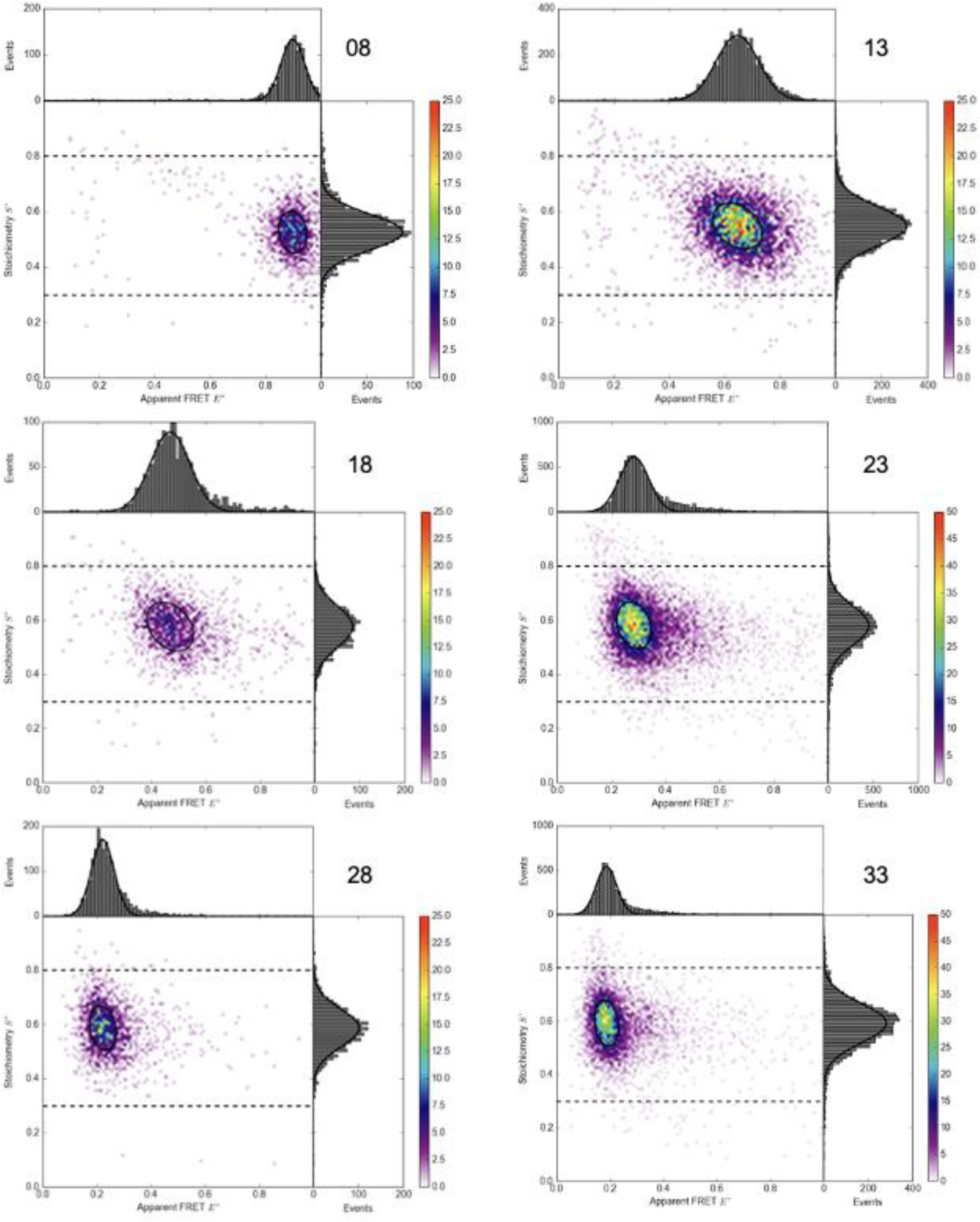
Two-dimensional ALEX-histograms showing setup-dependent uncorrected apparent E* vs S* values from single-molecule observation of dsDNA samples using the home-built confocal microscopy setup. The dsDNA ladder was labeled with Cy3B and Atto647N. The corresponding distances are 8 bp (top-left), 13 bp (top-right), 18 bp (middle-left), 23 bp (middle-right), 28 bp (lower-left) and 33 bp (lower-right).

**Supplementary Table 5:** Summary of fit results (2D Gaussian model) of smFRET data obtained with the home-built confocal microscopy for a Cy3 and Atto647N dsDNA ladder.

| bp | 8 | 13 | 18 | 23 | 28 | 33 |
| --- | --- | --- | --- | --- | --- | --- |
| $E^*$ | 0.91 | 0.70 | 0.50 | 0.32 | 0.27 | 0.23 |
| $\sigma_{E^*}$ | 0.04 | 0.08 | 0.09 | 0.08 | 0.07 | 0.06 |
| $S^*$ | 0.46 | 0.38 | 0.33 | 0.30 | 0.29 | 0.29 |
| $\sigma_{S^*}$ | 0.07 | 0.06 | 0.06 | 0.06 | 0.06 | 0.06 |

**Supplementary Figure 14:**
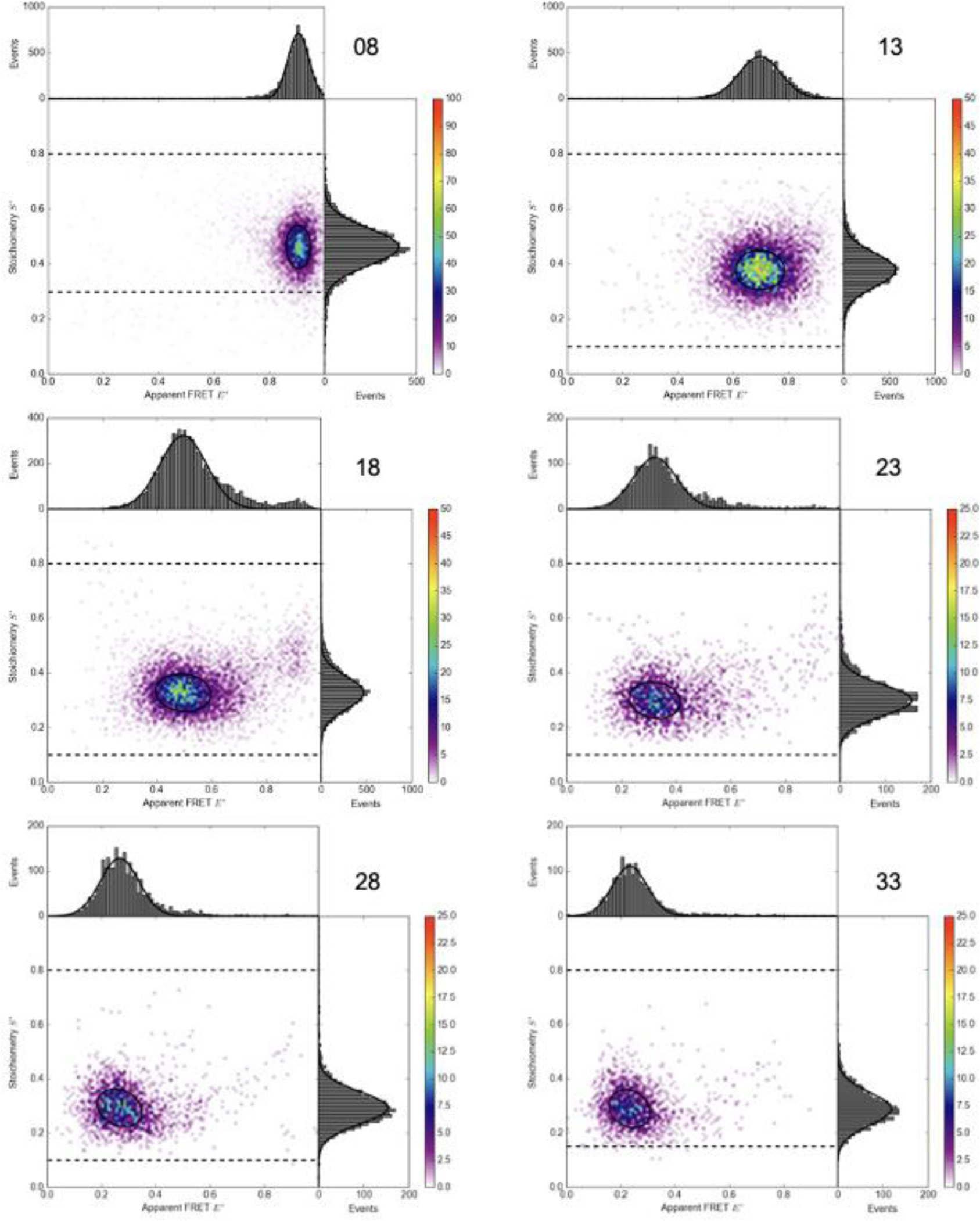
Two-dimensional ALEX-histograms showing setup-dependent uncorrected apparent E* vs S* values from single-molecule observation of dsDNA samples using the home-built confocal microscopy setup. The dsDNA ladder was labeled with Cy3 and Atto647N. The corresponding distances are 8 bp (top-left), 13 bp (top-right), 18 bp (middle-left), 23 bp (middle-right), 28 bp (lower-left) and 33 bp (lower-right).

**Supplementary Table 6:** Summary of fit results (2D Gaussian model) of smFRET data obtained with the home-built confocal microscopy for a Cy3B and Cy5 dsDNA ladder.

| bp | 8 | 13 | 18 | 23 | 28 | 33 |
| --- | --- | --- | --- | --- | --- | --- |
| $E^*$ | 0.93 | 0.67 | 0.46 | 0.27 | 0.20 | 0.18 |
| $\sigma_{E^*}$ | 0.04 | 0.09 | 0.07 | 0.06 | 0.04 | 0.04 |
| $S^*$ | 0.55 | 0.57 | 0.62 | 0.69 | 0.70 | 0.70 |
| $\sigma_{S^*}$ | 0.08 | 0.08 | 0.08 | 0.07 | 0.07 | 0.07 |

**Supplementary Figure 15:**
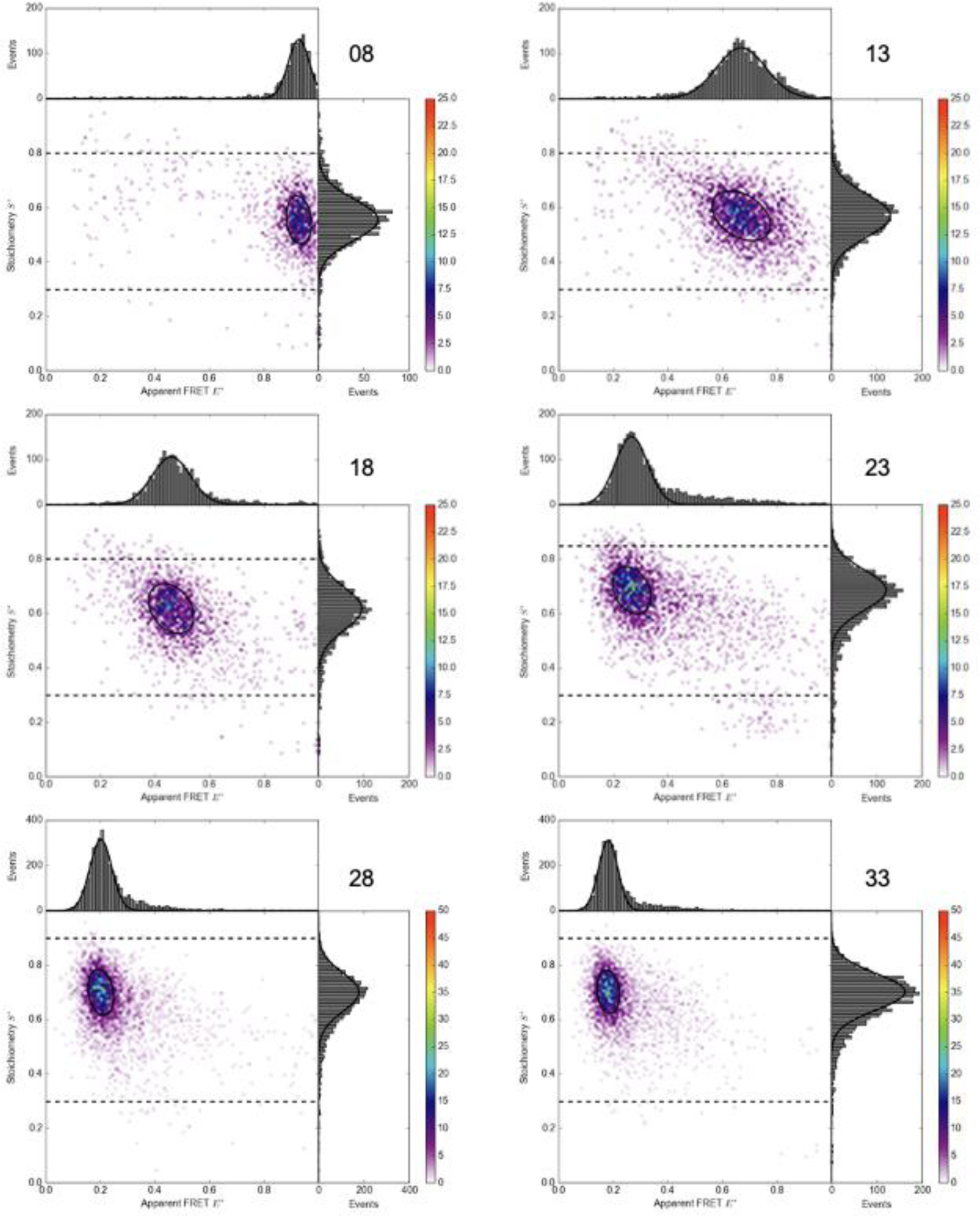
Two-dimensional ALEX-histograms showing setup-dependent uncorrected apparent E* vs S* values from single-molecule observation of dsDNA samples using the home-built confocal microscopy setup. The dsDNA ladder was labeled with Cy3B and Cy5. The corresponding distances are 8 bp (top-left), 13 bp (top-right), 18 bp (middle-left), 23 bp (middle-right), 28 bp (lower-left) and 33 bp (lower-right).

**Supplementary Table 7:** Accurate FRET analysis of DNA Cy3B-Atto647N measured with Brick-MIC. . Correction factors for leakage α, direct excitation δ, detection corrections β and γ and accurate FRET values and calculated distance.

| bp | 8 |  | 13 |  | 18 |  | 23 |  | 28 |  | 33 |  |
| --- | --- | --- | --- | --- | --- | --- | --- | --- | --- | --- | --- | --- |
| $\alpha$ | 0.081 | | 0.088 | | 0.082 | | 0.082 | | 0.081 | | 0.083 | |
| $\pm \Delta_{\alpha}$ | 0.00<br>4 | 0.00<br>4 | 0.01 | 0.02 | 0.00<br>6 | 0.00<br>5 | 0.00<br>1 | 0.00<br>1 | 0.00<br>6 | 0.00<br>5 | 0.00<br>5 | 0.00<br>7 |
| $\delta$ | 0.038 | | 0.041 | | 0.044 | | 0.044 | | 0.046 | | 0.045 | |
| $\pm \Delta_{\delta}$ | 0.00<br>4 | 0.00<br>8 | 0.00<br>2 | 0.00<br>3 | 0.00<br>4 | 0.00<br>2 | 0.00<br>2 | 0.00<br>4 | 0.00<br>2 | 0.00<br>2 | 0.00<br>2 | 0.00<br>3 |
| $\beta$ | 1.00 | | | | | | | | | | | |
| $\pm \Delta_{\beta}$ | 0.050 | | | | | | 0.03 | | | | | |
| $\gamma$ | 0.51 | | | | | | | | | | | |
| $\pm \Delta_{\gamma}$ | 0.11 | | | | | | 0.07 | | | | | |
| $S_{accMean}$ | 0.495 | | 0.503 | | 0.515 | | 0.54 | | 0.498 | | 0.48 | |
| $\pm \Delta_{Sacc}$ | 0.00<br>3 | 0.00<br>3 | 0.01<br>5 | 0.00<br>8 | 0.01<br>4 | 0.01<br>1 | 0.09 | 0.05 | 0.00<br>2 | 0.00<br>3 | 0.02 | 0.02 |
| $E_{accMean}$ | 0.902 | | 0.68 | | 0.470 | | 0.19 | | 0.122 | | 0.06 | |
| $\pm \Delta_{Eacc}$ | 0.00<br>7 | 0.00<br>9 | 0.05 | 0.04 | 0.00<br>9 | 0.00<br>5 | 0.04 | 0.07 | 0.00<br>9 | 0.00<br>8 | 0.03 | 0.02 |
| $R \text{ [\AA]}$ | 46.3 | | 59.1 | | 68.4 | | 86 | | 93.2 | | 108 | |
| $\pm \Delta_R \text{ [\AA]}$ | 0.1 | 0.7 | 0.5 | 2.2 | 0.2 | 0.4 | 7 | 3 | 0.2 | 1.3 | 6 | 8 |

**Supplementary Table 8:** Accurate FRET analysis of DNA Cy3B-Cy5 measured with Brick-MIC. . Correction factors for leakage α, direct excitation δ, detection corrections β and γ and accurate FRET and calculated distance values.

| bp | 8 |  | 13 |  | 18 |  | 23 |  | 28 |  | 33 |  |
| --- | --- | --- | --- | --- | --- | --- | --- | --- | --- | --- | --- | --- |
| $\alpha$ | 0.075 | | 0.077 | | 0.073 | | 0.078 | | 0.073 | | 0.073 | |
| $\pm \Delta_{\alpha}$ | 0.00<br>2 | 0.00<br>2 | 0.00<br>5 | 0.00<br>4 | 0.00<br>1 | 0.00<br>2 | 0.00<br>6 | 0.00<br>4 | 0.001<br>0 | 0.001<br>0 | 0.00<br>7 | 0.00<br>6 |
| $\delta$ | 0.05 | | 0.05 | | 0.081 | | 0.079 | | 0.087 | | 0.079 | |
| $\pm \Delta_{\delta}$ | 0.04 | 0.05 | 0.03 | 0.05 | 0.00<br>7 | 0.01<br>2 | 0.01<br>2 | 0.01<br>4 | 0.004 | 0.003 | 0.00<br>4 | 0.00<br>3 |
| $\beta$ | 0.90 | | | | | | | | | | | |
| $\pm \Delta_{\beta}$ | 0.07 | | | | | | 0.04 | | | | | |
| $\gamma$ | 0.32 | | | | | | | | | | | |
| $\pm \Delta_{\gamma}$ | 0.06 | | | | | | 0.08 | | | | | |
| $S_{\text{accMean}}$ | 0.50 | | 0.49 | | 0.50 | | 0.51 | | 0.53 | | 0.53 | |
| $\pm \Delta_{S_{\text{acc}}}$ | 0.01 | 0.01 | 0.01 | 0.01 | 0.01 | 0.02 | 0.02 | 0.02 | 0.01 | 0.02 | 0.01 | 0.01 |
| $E_{\text{accMean}}$ | 0.955 | | 0.763 | | 0.53 | | 0.272 | | 0.124 | | 0.02 | |
| $\pm \Delta_{E_{\text{acc}}}$ | 0.00<br>5 | 0.00<br>3 | 0.00<br>7 | 0.01<br>4 | 0.03 | 0.02 | 0.01<br>1 | 0.01<br>4 | 0.014 | 0.007 | 0.06 | 0.06 |
| $R \text{ [\AA]}$ | 44.3 | | 60.7 | | 72.5 | | 86.9 | | 102 | | 110 | |
| $\pm \Delta_R$<br>$[\text{\AA}]$ | 0.5 | 0.8 | 0.8 | 0.4 | 0.9 | 1.4 | 1.0 | 0.8 | 1 | 2 | 11 | 11 |

**Supplementary Table 9:**
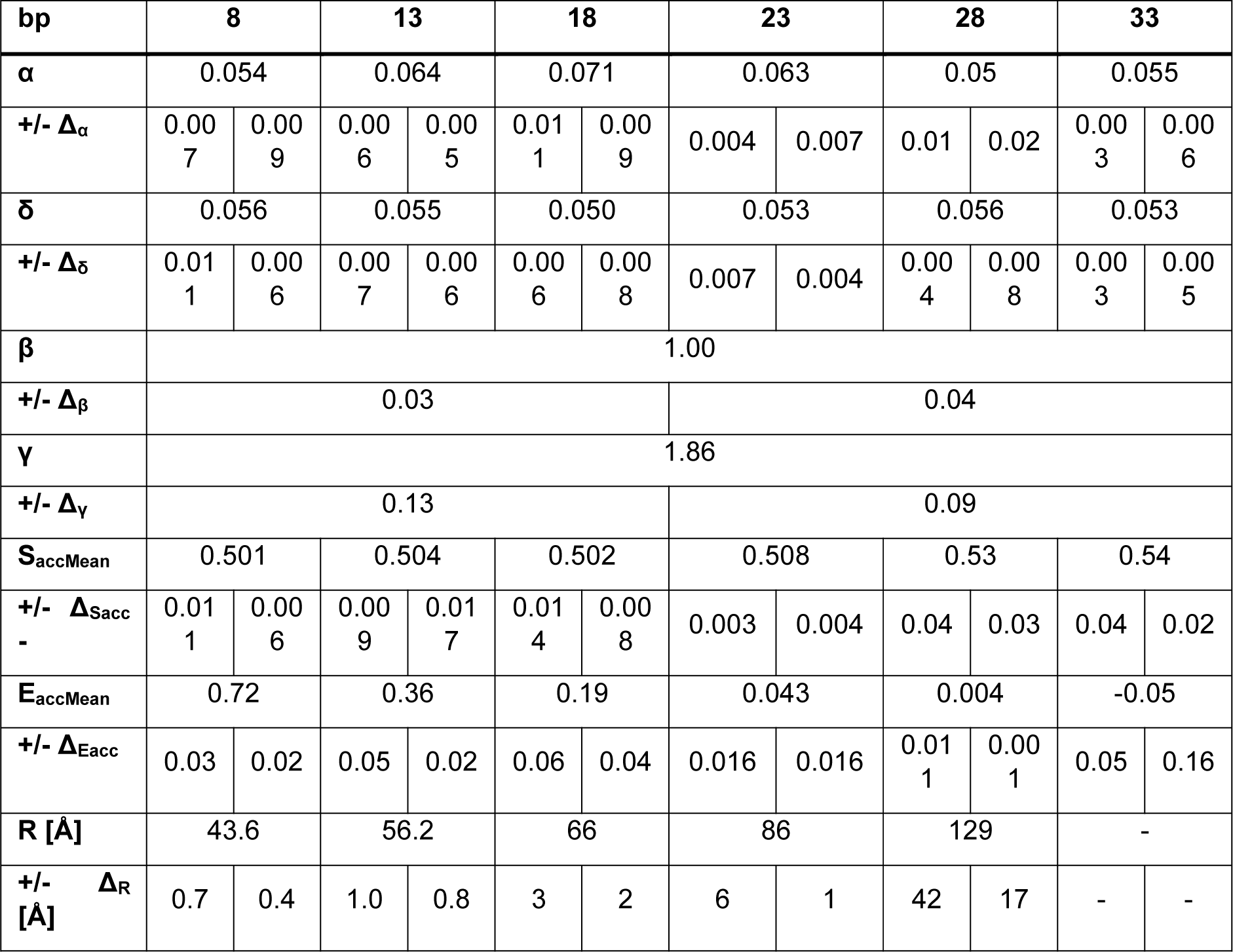
Accurate FRET analysis of DNA Cy3-Atto647N measured on with Brick-MIC. Correction factors for leakage α, direct excitation δ, detection corrections β and γ and accurate FRET values and calculated distance.

**Supplementary Table 10:**
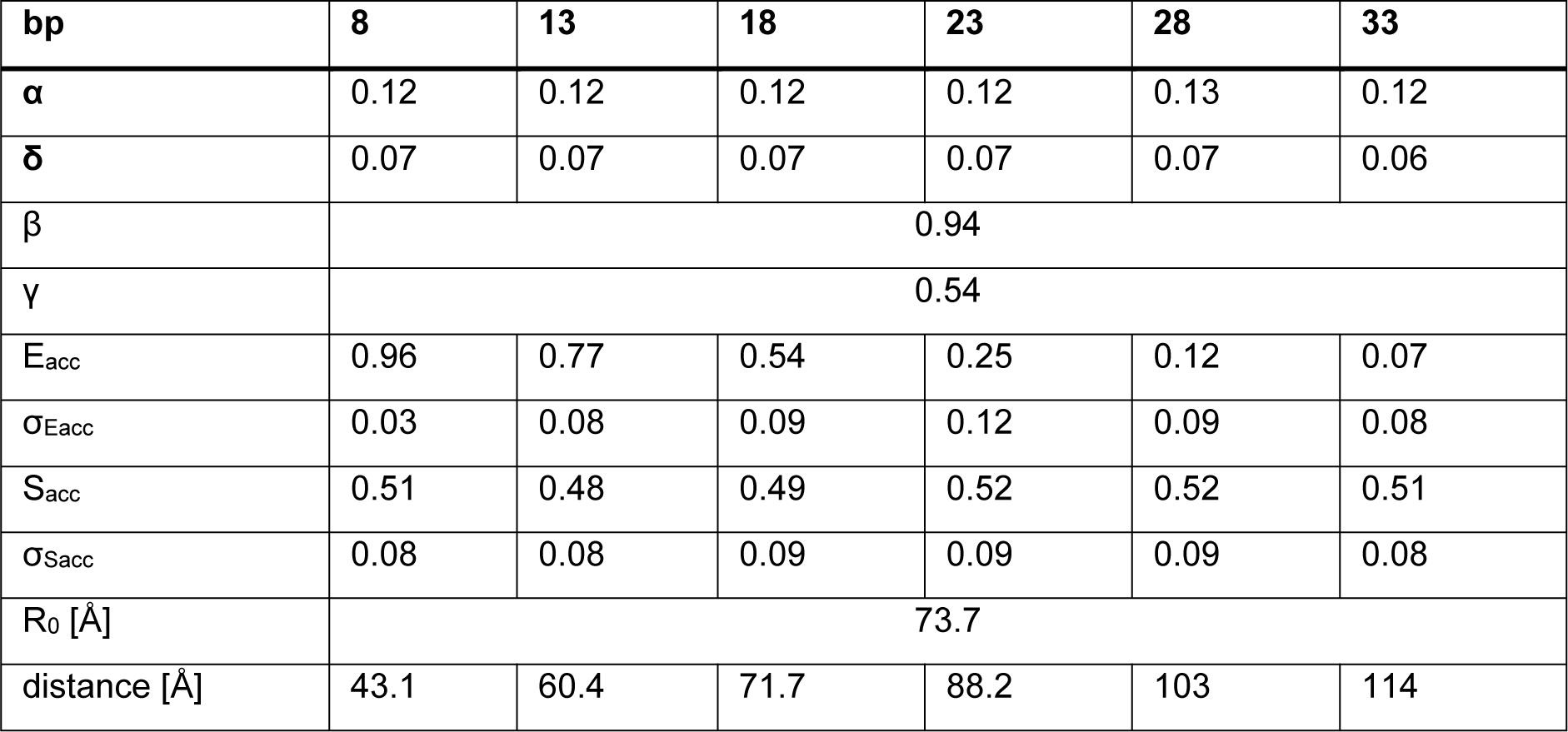
Accurate FRET analysis of DNA Cy3B-Cy5 measured on a home-built confocal microscope. . Correction factors for leakage α, direct excitation δ, detection corrections β and γ and accurate FRET values and calculated distances.

| bp | 8 | 13 | 18 | 23 | 28 | 33 |
| --- | --- | --- | --- | --- | --- | --- |
| $\alpha$ | 0.12 | 0.12 | 0.12 | 0.12 | 0.13 | 0.12 |
| $\delta$ | 0.07 | 0.07 | 0.07 | 0.07 | 0.07 | 0.06 |
| $\beta$ | 0.94 | | | | | |
| $\gamma$ | 0.54 | | | | | |
| $E_{acc}$ | 0.96 | 0.77 | 0.54 | 0.25 | 0.12 | 0.07 |
| $\sigma_{Eacc}$ | 0.03 | 0.08 | 0.09 | 0.12 | 0.09 | 0.08 |
| $S_{acc}$ | 0.51 | 0.48 | 0.49 | 0.52 | 0.52 | 0.51 |
| $\sigma_{Sacc}$ | 0.08 | 0.08 | 0.09 | 0.09 | 0.09 | 0.08 |
| $R_0$ [Å] | 73.7 | | | | | |
| distance [Å] | 43.1 | 60.4 | 71.7 | 88.2 | 103 | 114 |

**Supplementary Table 11:** Accurate FRET analysis of DNA Cy3B-Atto647N measured on a standard confocal microscope. . Correction factors for leakage α, direct excitation δ, detection corrections β and γ and accurate FRET values and calculated distances.

| bp | 8 | 13 | 18 | 23 | 28 | 33 |
| --- | --- | --- | --- | --- | --- | --- |
| $\alpha$ | 0.12 | 0.12 | 0.12 | 0.12 | 0.12 | 0.12 |
| $\delta$ | 0.04 | 0.02 | 0.02 | 0.04 | 0.04 | 0.04 |
| $\beta$ | 0.95 | | | | | |
| $\gamma$ | 0.82 | | | | | |
| $E_{acc}$ | 0.91 | 0.67 | 0.46 | 0.21 | 0.12 | 0.06 |
| $\sigma_{Eacc}$ | 0.04 | 0.08 | 0.09 | 0.08 | 0.06 | 0.06 |
| $S_{acc}$ | 0.50 | 0.50 | 0.51 | 0.50 | 0.50 | 0.50 |
| $\sigma_{Sacc}$ | 0.07 | 0.07 | 0.08 | 0.08 | 0.07 | 0.08 |
| $R_0$ [Å] | 67.0 | | | | | |
| distance [Å] | 45.2 | 59.7 | 68.8 | 83.4 | 93.3 | 105 |

**Supplementary Table 12:** Accurate FRET analysis of DNA Cy3-Atto647N measured on a standard confocal microscope. . Correction factors for leakage α, direct excitation δ, detection corrections β and γ and accurate FRET values and calculated distances.

| bp | 8 | 13 | 18 | 23 | 28 | 33 |
| --- | --- | --- | --- | --- | --- | --- |
| $\alpha$ | 0.09 | 0.10 | 0.12 | 0.10 | 0.09 | 0.10 |
| $\delta$ | 0.04 | 0.04 | 0.04 | 0.04 | 0.04 | 0.04 |
| $\beta$ | 1.09 | | | | | |
| $\gamma$ | 2.97 | | | | | |
| $E_{acc}$ | 0.77 | 0.39 | 0.18 | 0.06 | 0.03 | -0.04 |
| $\sigma_{Eacc}$ | 0.09 | 0.09 | 0.06 | 0.05 | 0.04 | 0.06 |
| $S_{acc}$ | 0.51 | 0.50 | 0.50 | 0.51 | 0.50 | 0.51 |
| $\sigma_{Sacc}$ | 0.07 | 0.07 | 0.07 | 0.07 | 0.08 | 0.09 |
| $R_0$ [Å] | 51.1 | | | | | |
| distance [Å] | 41.9 | 54.9 | 65.5 | 80.4 | 91.8 | - |

**Supplementary Table 13:** Comparison of correction factors β and γ of the BRICK-MIC and a standard confocal microscope.

|  | DNA-Cy3B-Atto647N |  | DNA-Cy3B-Cy5 |  | DNA-Cy3-Atto647N |  |
| --- | --- | --- | --- | --- | --- | --- |
|  | Standard Confocal | Brick-MIC | Standard Confocal | Brick-MIC | Standard Confocal | Brick-MIC |
| $\beta$ | 0.95 | 1.00 | 0.94 | 0.90 | 1.09 | 1.00 |
| $\gamma$ | 0.82 | 0.51 | 0.54 | 0.32 | 2.97 | 1.86 |

**Supplementary Table 14:** E*-S*-fit results of MalE (T36C-S352) labelled with Alexa555-Alexa647 measured with Brick-MIC and standard confocal microscope.

|  | Brick-MIC |  | Standard confocal |  |
| --- | --- | --- | --- | --- |
| $C_{maltose}$ [mM] | 0 | 1 | 0 | 1 |
| $E^*$ | 0.57 | 0.76 | 0.68 | 0.84 |
| $\sigma_{E^*}$ | 0.08 | 0.07 | 0.07 | 0.05 |
| $S^*$ | 0.49 | 0.53 | 0.46 | 0.51 |
| $\sigma_{S^*}$ | 0.06 | 0.06 | 0.06 | 0.06 |

**Supplementary Table 15:** Accurate FRET analysis of MalE (T36C-S352) labelled with Alexa555-Alexa647, measured with Brick-MIC (left) and a home-built confocal microscope (right). Using the correction factors α, δ, β and γ accurate FRET values and the distances for the opened and closed conformation, apo and holo state, respectively, of MalE (T36C-S352) were calculated. The Förster radius for Alexa555 and Alexa647 R0 = 51 Å. *DD and AA only E- and S-values were determined from data sets with a significant DD- and AA-only population with the same protein and fluorophore.

|  | Brick-MIC, ALEX |  |  |  | Standard confocal microscope |  |
| --- | --- | --- | --- | --- | --- | --- |
| C<sub>maltose</sub>[mM] | 0 |  | 1 |  | 0 | 1 |
| α | 0.060 |  | 0.0578 |  | 0.10* |  |
| +/- Δ<sub>α</sub> | 0.004 | 0.003 | 0.0003 | 0.0003 | - | - |
| δ | 0.082 |  | 0.092 |  | 0.07* |  |
| +/- Δ<sub>δ</sub> | 0.011 | 0.013 | 0.007 | 0.008 | - | - |
| β | 0.83 |  |  |  | 0.82 |  |
| +/- Δ<sub>β</sub> | 0.04 |  | 0.08 |  | - | - |
| γ | 1.82 |  |  |  | 2.34 |  |
| +/- Δ<sub>γ</sub> | 0.15 |  | 0.09 |  | - | - |
| S<sub>accMean</sub> | 0.493 |  | 0.499 |  | 0.50 | 0.49 |
| +/- Δ<sub>Sacc</sub> - | 0.004 | 0.005 | 0.003 | 0.003 | - | - |
| E<sub>accMean</sub> | 0.36 |  | 0.62 |  | 0.39 | 0.66 |
| +/- Δ<sub>Eacc</sub> | 0.01 | 0.03 | 0.03 | 0.04 | - | - |
| R [Å] | 56.2 |  | 47.2 |  | 54.9 | 45.7 |
| +/- Δ<sub>R</sub> [Å] | 0.5 | 0.5 | 0.4 | 1.1 | - | - |

**Supplementary Table 16:** Comparison of measured and simulated distances of the conformational states of MalE (T36C-S352) labelled with Alexa555-Alexa647. All distances in the table provided in Å.

| Conformational state of MalE1 | Apo | Holo |
| --- | --- | --- |
| AV calculations | 57.5 | 46.3 |
| Brick-MIC | 56.2 | 47.2 |
| Standard confocal | 54.9 | 45.7 |

**Supplementary Fig. 16:**
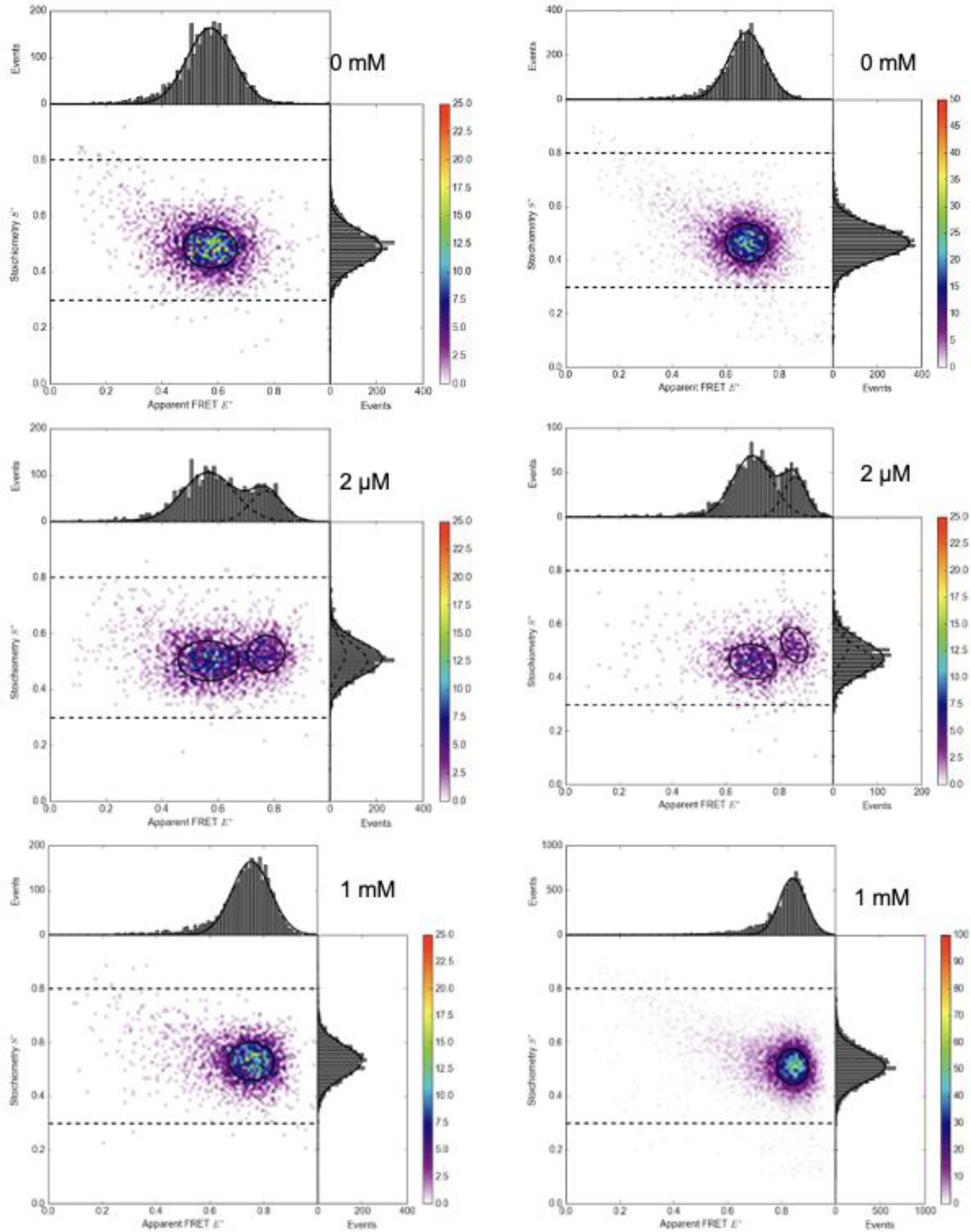
ALEX-histograms of MalE (T36C-S352) labelled with Alexa555-Alexa647 and studied by the APD variant of Brick-MIC (left) and a home-built confocal microscope (right). Data are shown as two-dimensional histograms with setup-dependent uncorrected apparent E* vs S* values from single-molecule observation of multiple bursts. Experimental data for maltose concentrations 0 mM, 2 µM and 1 mM, reflecting the apo, KD and holo states, respectively.

**Supplementary Fig. 17:**
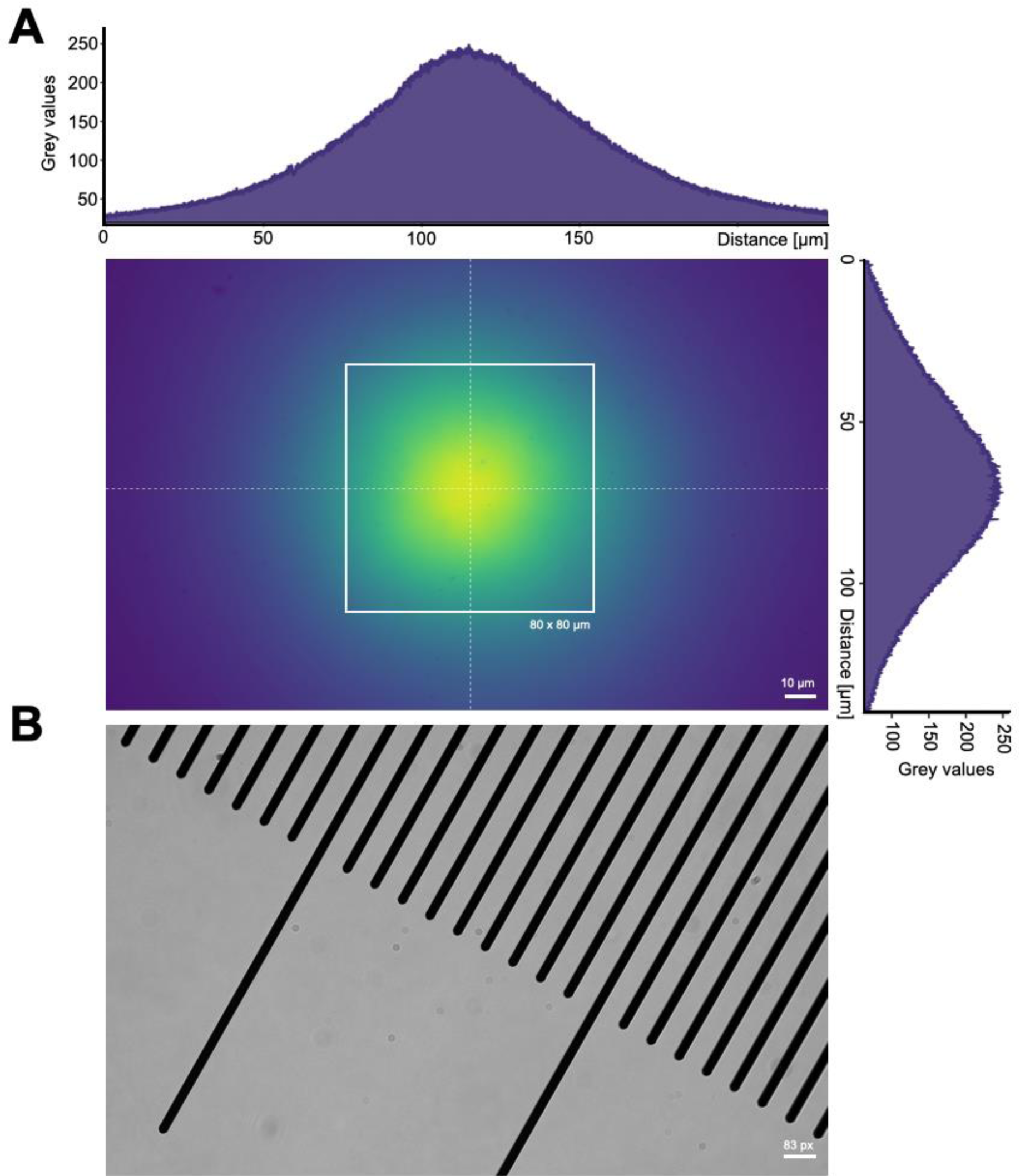
Characterizing the field of view and laser intensity distribution of the epi-fluorescence variant of Brick-MIC. Panel A) shows the Gaussian excitation profile acquired by imaging a red fluorescent slide (FSK5, Thorlabs) excited at 640 nm wavelength with 1 mW laser power. The dotted lines depicts the location of the x/y intensity profile. The rectangle indicates the field of view used for STORM and PAINT experiments; these dimensions were selected as one standard deviation from the Gaussian distribution. Panel B shows a transmission light microscopy image of a micrometer calibration slide with 10 μm interspaced markings.

**Supplementary Fig. 18:**
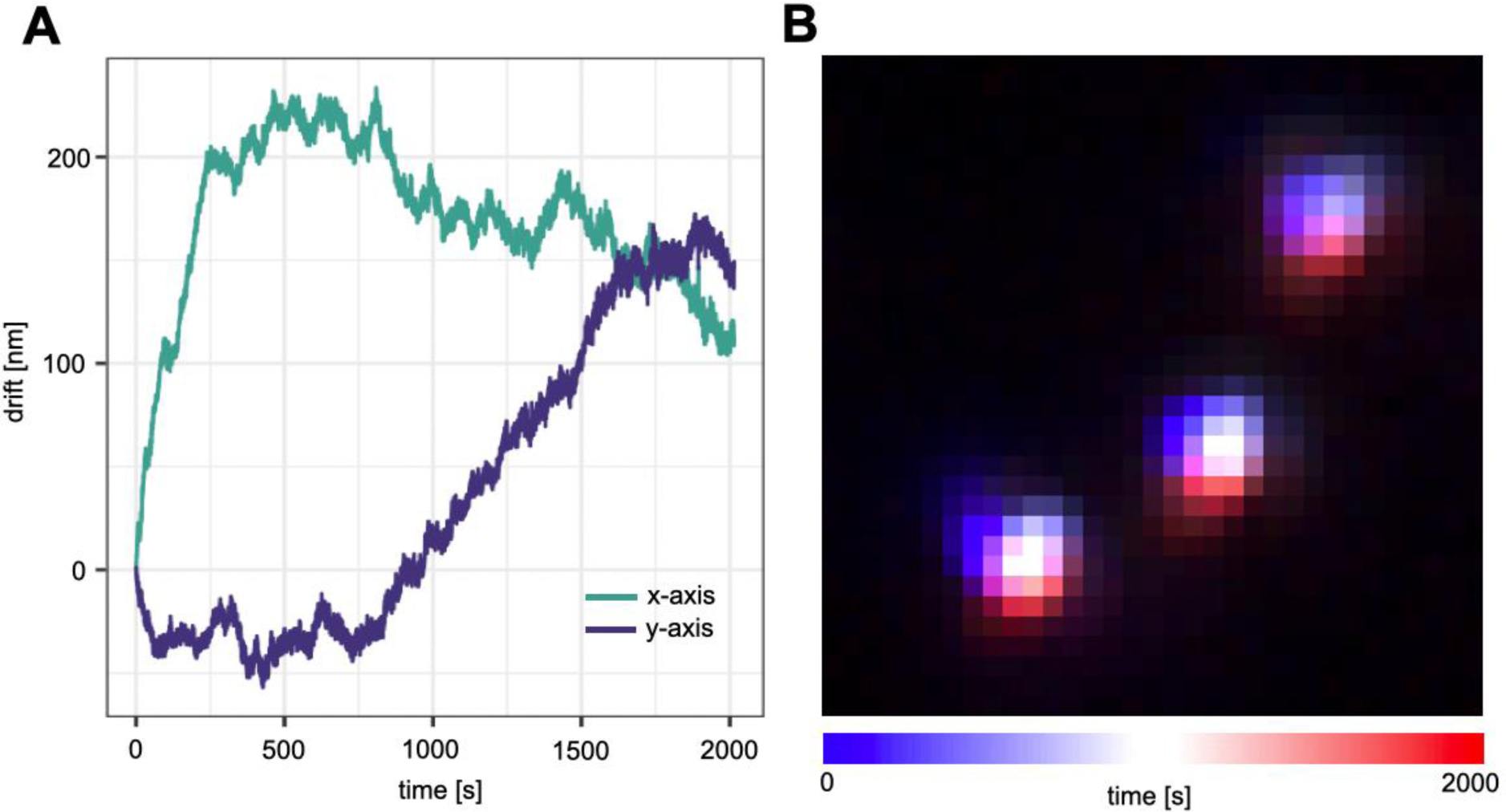
Quantifying long-term lateral drift. Panel A depicts the x-y drift of 100-nm-sized Tetraspeck beads (image shown in panel B) over a time interval of 2000 s, where the initial location is shown in blue and later locations in red.

**Supplementary Fig. 19:**
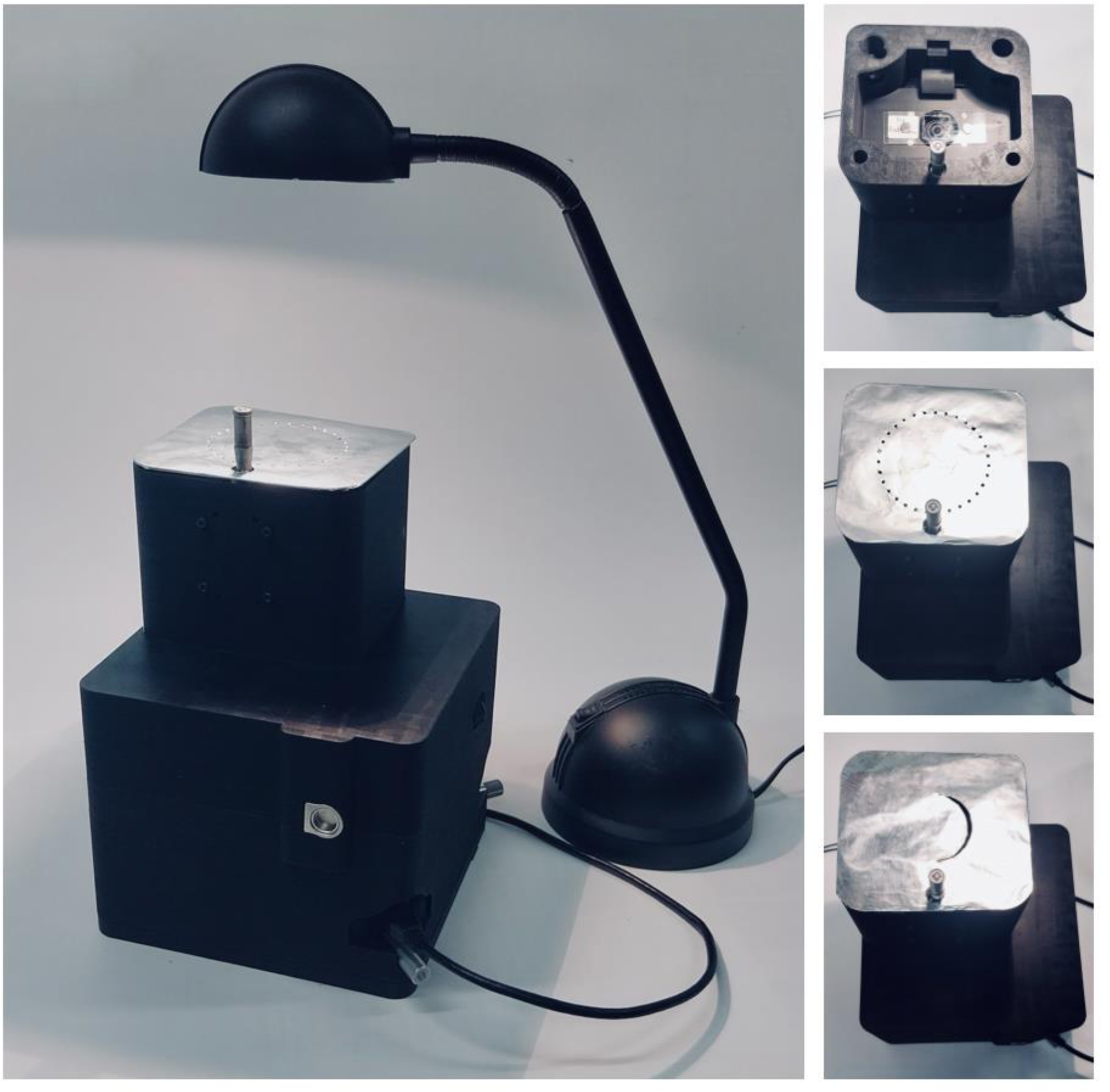
Transmission light microscopy using Brick-MIC. The left panel shows the Brick-MIC video modality with an LED desktop lamp used as light source, which was positioned right on top of the sample holder. The sample holder is covered with aluminum foil with a punctured circular pattern for darkfield imaging (middle right panel). The upper right panel shows an uncovered sample holder used for bright field imaging. The lower right panel shows the aluminum covering used for oblique illumination, where the foil has a punctured crescent moon shape.

## Notes

https://zenodo.org/records/10441063

### Summary of Updates

Main text was adapted according to referee comments, all figures were updated and a new version of the data acqusition and analysis software are available. Details are found in the deposited pdf.

